# Translatome analysis reveals microglia and astrocytes to be distinct regulators of inflammation in the hyperacute and acute phases after stroke

**DOI:** 10.1101/2023.02.14.520351

**Authors:** Victoria G. Hernandez, Kendra J. Lechtenberg, Todd C. Peterson, Li Zhu, Tawaun A. Lucas, Justice O. Owah, Alanna I. Dorsey, Andrew J. Gentles, Marion S. Buckwalter

**Author notes:** **Corresponding author:** Marion Buckwalter. Shared first authors (Victoria G. Hernandez, Kendra J. Lechtenberg). **Author Information:** Victoria G. Hernandez, Kendra J. Lechtenberg, Todd C. Peterson, Li Zhu, Tawaun A. Lucas, Justice O. Owah, Alanna I. Dorsey, Andrew J. Gentles. **Data availability statement:** Raw translatome datasets generated from microglia, astrocytes, and whole brain cortex are being uploaded to GEO (Accession # GSE225110) as fastq files. To improve visualization of the complete processed data set, a web application was developed; this can be found at https://buckwalterlab.shinyapps.io/AstrocyteMicrogliaRiboTag/.

## Abstract

Neuroinflammation is a hallmark of ischemic stroke, which is a leading cause of death and long-term disability. Understanding the exact cellular signaling pathways that initiate and propagate neuroinflammation after stroke will be critical for developing immunomodulatory stroke therapies. In particular, the precise mechanisms of inflammatory signaling in the clinically relevant hyperacute period, hours after stroke, have not been elucidated. We used the RiboTag technique to obtain astrocyte and microglia-derived mRNA transcripts in a hyperacute (4 hours) and acute (3 days) period after stroke, as these two cell types are key modulators of acute neuroinflammation. Microglia initiated a rapid response to stroke at 4 hours by adopting an inflammatory profile associated with the recruitment of immune cells. The hyperacute astrocyte profile was marked by stress response genes and transcription factors, such as *Fos* and *Jun*, involved in pro-inflammatory pathways such as TNF-α. By 3 days, microglia shift to a proliferative state and astrocytes strengthen their inflammatory response. The astrocyte pro-inflammatory response at 3 days is partially driven by the upregulation of the transcription factors *C/EBPβ, Spi1*, and *Rel*, which comprise 25% of upregulated transcription factor-target interactions. Surprisingly, few sex differences across all groups were observed. Expression and log_2_ fold data for all sequenced genes are available on a user-friendly website for researchers to examine gene changes and generate hypotheses for stroke targets. Taken together our data comprehensively describe the astrocyte and microglia-specific translatome response in the hyperacute and acute period after stroke and identify pathways critical for initiating neuroinflammation.

## Introduction

Stroke is a leading cause of death and long-term disability worldwide (Virani et al., 2021). Despite its prevalence, existing treatments for ischemic stroke are limited to restoration of blood flow to the affected brain tissue by using tissue plasminogen activator (tPA) or mechanical methods to remove blockages. This serves to mitigate the extent of initial cell death; however, these interventions are only effective within a narrow time window (<24 hrs), and other pharmacologic therapies are extremely limited. Neuroinflammation after stroke is a promising therapeutic target because it leads to increased pathophysiology, neurodegeneration and poorer outcomes in the days to months following the initial insult. Unfortunately, clinical trials targeting the overall curtailment of inflammation have been unsuccessful in improving stroke outcomes (Moretti et al., 2015; Iadecola et al., 2020; Stuckey et al., 2021).

To effectively modulate inflammation after stroke to improve outcomes, it is critical to define immune changes in the hyperacute phase that affect the ensuing cascade of molecular processes including inflammation and its resolution and its repair and regeneration. Resolution of inflammation depends on rapid clearance of early damage signals, upregulation of anti-inflammatory cytokines, and removal of dead cells (Shichita et al., 2017), yet the molecular signature of the hyperacute inflammatory response that initiates neuroinflammation after stroke is not known. We do know that this response is largely initiated by brain resident glial cells. Many of the hallmarks of neuroinflammation following stroke–release of pro-inflammatory cytokines, recruitment of immune cells, formation of the glial scar, and modulation of the blood brain barrier (BBB)–are largely initiated by microglia and astrocytes during this hyperacute period. Inhibition or depletion of microglia before stroke results in larger strokes and worse functional outcomes (Lalancette-Hébert et al., 2007; Szalay et al., 2016). However, prolonged or exaggerated microglial activation after stroke is also believed to be detrimental (Iadecola and Anrather, 2011). Astrocytes play a similarly complex role after stroke, both amplifying inflammation (Rakers et al., 2019) and forming a glial scar that can be protective and promote axonal regeneration (Faulkner et al., 2004; Anderson et al., 2016, 2018). The potentially contradictory roles of microglia and astrocytes in neuroprotection after ischemic injury underscore the complex nature of inflammation and how it might divergently impact recovery at different stages following stroke.

For these reasons, we sought to accurately characterize the microglial and astrocytic translatome in the hyperacute period after stroke. We achieved this by using a ribosomal pulldown technique to isolate microglia or astrocyte-enriched RNA (Sanz et al., 2009). This strategy eliminates the need to isolate cells by sorting, which can introduce early responses that are similar enough to early injury responses to confound effects (Haimon et al., 2018). The ribosomal pulldown technique also allows for rapid isolation of RNA at a precise time point. We used immunoprecipitation in Microglia RiboTag and Astrocyte RiboTag mice to rapidly isolate actively translating mRNA from cortical tissue. We selected 4 hours as a hyperacute time point after stroke as it is a clinically accessible yet very early window in terms of defining the initiation of inflammation, and 3 days as an acute time point when inflammation is well underway, and may serve as a second treatment window for patients unable to receive hyperacute interventions. The distal middle cerebral artery occlusion model (dMCAO) of ischemic stroke was chosen for this study to model ischemia without reperfusion. The dMCAO model is advantageous due to its accessibility and reproducibility: it results in predominantly cortical strokes and has low variability in stroke size and location (Tamura et al., 1981; Doyle et al., 2012). We analyzed changes in microglial and astrocyte gene expression at a hyperacute and acute time point after stroke and applied differential expression and pathway analysis to identify key features of the microglial and astrocyte response at each of these post-stroke time points. Finally, we constructed a user-friendly web resource for the community to use to investigate their genes of interest after stroke.

## Methods

### 2.1 Animals

All animal use was in accordance with protocols approved by the Stanford University Institutional Animal Care and Use Committee. Cx3cr1^CreER^ mice (B6.129P2(Cg)-*Cx3cr1*^*tm2*.*1(cre/ERT2)Litt*^/WganJ, JAX:021160), Aldh1l1^CreER^ (B6N.FVB-Tg(Aldh1l1-cre/ERT2)1Khakh/J, JAX:029655) and Rpl22^HA^ (B6.129(Cg)-*Rpl22*^*tm1*.*1Psam*^/SjJ, JAX:029977) were purchased from The Jackson Laboratories (Bar Harbor, ME). Mice were housed in a temperature-controlled 12-hour light-dark alternating facility, with *ad libitum* access to food and water. All experiments were performed with 10 to 12-week-old male and female mice. To generate mice with conditional expression of a hemagglutinin (HA) tag on the Rpl22 ribosomal subunit in microglia and astrocytes, we bred Cx3cr1^CreER^ and Aldh1l1^CreER^ animals with Rpl22^HA+/+^ animals, respectively. Experimental animals, *Cx3cr1*^*CreER2-IRES-eYFP/+*^*;Rpl22*^*HA+/+*^ (Microglia RiboTag) and *Aldh1l1*^*CreER2/+*^*;Rpl22*^*HA+/+*^ (Astrocyte RiboTag) were homozygous for the *Rpl22*^HA^ allele and heterozygous for the CreER knock-in allele. *Cx3cr1*^*CreER2-IRES-eYFP/+*^*;Rpl22*^*HA+/+*^ animals were treated with 200 mg/kg tamoxifen in corn oil via oral gavage on three consecutive days to induce recombination and expression of the hemagglutinin tag. Stroke surgeries were performed 30 days after the last tamoxifen dose. *Aldh1l1*^*CreER2/+*^*;Rpl22*^*HA+/+*^ animals were treated with 80 mg/kg tamoxifen (Srinivasan et al., 2016) in corn oil via oral gavage on five consecutive days to induce recombination and expression of the hemagglutinin tag. Stroke surgeries were performed 8 days after the last tamoxifen dose to ensure tamoxifen and its metabolites were completely degraded (Valny et al., 2016).

### 2.2 Distal middle cerebral artery occlusion stroke model

Stroke was induced using distal middle cerebral artery occlusion (dMCAO) which has been previously described in detail (Tamura et al., 1981; Doyle et al., 2012). Briefly, animals were anesthetized with 2% Isoflurane in 2 L/min 100% oxygen and an incision was made along the animal’s skull, first through the skin and then through the temporalis muscle. The middle cerebral artery was identified, a craniotomy was drilled, dura was removed, and the artery was cauterized. Sham mice were subject to the same surgical procedure as stroke mice, except that the artery was not cauterized. Animals were maintained at 37° C both during surgery and recovery using a feedback-controlled heating blanket. Following surgery, incisions were sealed using Surgi-lock tissue adhesive (Meridian, Allegan, MI). Mice were concurrently injected with 25 mg/kg cefazolin (VWR #89149-888) and 0.5 mg/kg of buprenorphine SR (Zoopharm, Windsor, CO) to prevent infection and for pain management, respectively. Animals were monitored until ambulatory and for infection at the surgical site post-operation.

### 2.3 Immunohistochemistry

Mice were deeply anesthetized with either chloral hydrate or a ketamine/xylazine cocktail and perfused with 0.9% heparinized saline. Brains were collected and drop-fixed in 4% PFA in phosphate buffer for 24 hours at 4°C, then preserved in 30% sucrose in PBS. A freezing microtome (Microm HM430) was used to collect 40 μm thick coronal brain sections sequentially into 12 tubes. Brain sections were stored in cryoprotectant medium (30% glycerin, 30% ethylene glycol, and 40% 0.5 M sodium phosphate buffer) at −20°C until processing. Standard immunohistochemistry procedures were used to stain free floating sections. Briefly, sections were blocked with 3% donkey serum (Millipore, #S30-100mL) for one hour, then incubated for 15 minutes in each of streptavidin and biotin blocking solutions (Vector Laboratories, SP-2002). Tissue was then incubated at 4°C overnight in primary antibody: anti-Iba1 (rabbit, 1:1000, Wako 019-19741), biotinylated anti-HA (mouse, 1:500, BioLegend 901505), anti-CD68 (rat, 1:1000, BioRad MCA1957S), or anti-GFAP (rat, 1:500, Invitrogen 13-0300). GFAP, Iba1, and HA were labeled with fluorescent secondary antibodies (donkey anti-rat IgG, 1:200; donkey anti-rabbit IgG, 1:200, Thermo-Fisher A-31573; anti-streptavidin IgG, 1:200, Invitrogen Molecular Probes S-32355), mounted onto glass slides, and cover slipped using Vectashield HardSet Mounting Medium (Vector Laboratories, H-1400). For CD68 staining, tissue was incubated for one hour in secondary antibody (rabbit anti-rat IgG, 1:500, Vector Laboratories BA-4001) treated with Avidin-Biotin Complex solution (Vector, #PK-6100) for one hour, and treated for 5 minutes with filtered DAB (Sigma, #D5905) solution. Finally, sections were mounted onto glass slides, air-dried overnight, and then cover slipped with Entellan (Electron Microscopy Sciences 14800).

### 2.4 Image acquisition and quantification of HA colocalization

Z-stacks were taken at 40X magnification using a Leica confocal microscope. Three z-stacks per brain section were taken in the peri-infarct region dorsal, lateral, and ventral to the stroke lesion, or in equivalent locations in sham brain sections. Five sections were imaged per mouse. Quantification of colocalization between HA, Iba1, Cx3cr1-eYFP, and GFAP was performed using ImageJ software, and counts were averaged across all images for each animal. Then, we calculated the percentage of HA/Iba1/Cx3cr1 triple-positive cells and HA/GFAP double-positive cells by dividing by the total number of HA+ cells to quantify specificity of HA expression. We calculated the percentage of Cx3cr1-eYFP/HA and GFAP/HA double-positive cells out of the total number of GFAP+ cells to quantify efficiency of recombination. All imaging and quantification was performed by an experimenter blinded to the experimental group.

### 2.5 Ribosome immunoprecipitation and RNA extraction

Mice were deeply anesthetized with chloral hydrate or a ketamine/xylazine cocktail and sacrificed at either a hyperacute (4 hours after stroke) or acute (3 days after stroke) time point. Peri-infarct cortical tissue and stroke core was rapidly dissected, flash-frozen using liquid nitrogen, and stored at −80°C to prevent RNA degradation. Peri-infarct cortex was defined as the area of cortical tissue within a 2.5 mm radius of the stroke core at the time of dissection. Immunoprecipitation was performed as previously described (Sanz et al., 2013); (Sanz et al., 2009). Briefly, brain tissue was homogenized at 3% weight per volume using a Dounce homogenizer in buffer (50 mM Tris, pH 7.5, 100 mM KCl, 12 mM MgCl_2_, and 1% NP-40) containing 100 μg/mL cycloheximide (Sigma), 1 mM DTT, 200 U/mL RNAsin (Promega), 1 mg/mL heparin, 1X Protease inhibitors (Sigma) and 1 mM DTT. Homogenates were centrifuged for 10 minutes at 10,000 rpm and 4°C. A portion of the supernatant was immediately stored at −80°C and subsequently served as the “input” for each sample. The remainder of the supernatant was incubated with anti-HA antibody (1:100, BioLegend 901502) on a gentle rotator for 4 hours at 4°C, followed by addition of 200 μL of Dynabeads Protein G (Invitrogen 10004D) and overnight incubation at 4°C with gentle rotating.

Tubes were placed on a magnetic stand to allow removal of the supernatant, and then polysome-bead-antibody complexes were washed 5X at 4°C with high salt buffer (50 mM Tris, pH 7.5, 300 mM KCl, 12 mM MgCl_2_, 1% NP-40, 1 mM DTT, and 100 μg/mL cycloheximide). Buffer RLT with β-mercaptoethanol (Qiagen RNeasy Micro Kit) was added to the samples immediately after the last wash and tubes were vortexed to release the polysomes from the beads. Tubes were placed back on the magnetic stand, and supernatant containing the polysomes was stored at −80°C until RNA extraction. These samples were subsequently designated as immunoprecipitated, or “IP”. For both IP and input samples, genomic DNA was removed, and RNA was extracted according to manufacturer’s protocols using an RNeasy Micro Kit (Qiagen) for IP and RNeasy Mini Kit (Qiagen) for input samples.

### 2.6 Library preparation and RNA sequencing

RNA integrity and concentration was measured by BioAnalyzer (Eukaryote RNA Pico, Agilent). Samples with an RNA integrity number (RIN) > 6 were used for RNA-sequencing (average RIN = 8.5). For microglia samples, polyA-enrichment of 8 ng of total RNA from each replicate was used to prepare a cDNA library using the Smart-Seq2 method (Picelli et al., 2014). Libraries were modified for RNA-sequencing using the Nextera XT DNA Sample Preparation Kit (Illumina) with 200 pg of cDNA as starting material. Library quality was assessed by Qubit fluorometer (high sensitivity dsDNA, ThermoFisher), BioAnalyzer (high sensitivity DNA, Agilent), and TapeStation (high sensitivity DNA, Agilent). Sequencing was performed on the Illumina HiSeq 4000 platform to obtain 150 bp paired-end reads. The astrocyte samples were sent to Azenta (formerly GeneWiz) for library preparation, using the NEBNext Ultra RNA Library Preparation Kit with polyA selection. Sequencing was performed on the Illumina HiSeq 4000 platform to obtain 150 bp paired-end reads.

### 2.7 RNAseq data processing and analysis

Approximately 12–36 million 150□bp reads were obtained from all microglia and astrocyte samples. These reads were trimmed of adapter sequences, low quality bases, and very short reads using trimGalore! (v0.4.5), a wrapper script which utilizes cutadapt (Martin, 2011) and FastQC (v0.11.7). Remaining reads were aligned to the mouse mm10 genome (GRCm38.p6) using the STAR aligner (Dobin et al., 2013) (v2.7.1), which utilizes an algorithm that minimizes alignment time, mapping errors, and alignment biases. Transcript abundances were then annotated and quantified using the RSEM software package (Li and Dewey, 2011) (v1.3.1). Differential gene expression analysis was carried out using the DESeq2 (Love et al., 2014) (v1.26) package within the R environment. The “apeglm” method was used to shrink log fold change calculations (Zhu et al., 2019). A significance cutoff was set using the adjusted *p*-value < 0.05 and |log_2_fold change| > 1. We eliminated some samples based on principal component analysis of the top 500 expressed genes. Gene ontology analysis was performed using DAVID (version 6.8) to identify biological processes and KEGG pathways enriched after stroke (Huang et al., 2009a; b). Four microglia samples presented as extreme outliers, and earlier experimental notes indicated that these particular brain samples appeared abnormal at the time of dissection (e.g. a. large hemorrhage in a sham brain, no visible stroke in a “stroke” brain). Four sequenced 4 hour sham astrocyte samples clustered with the 4 hour stroke astrocyte samples. To address this, we repeated all groups. In the new cohort, 4 hour sham samples clustered with the other 4 hour sham samples and the other groups clustered as expected. All were included in the analysis except the 4 mis-clustered sham samples from cohort 1. Transcripts with low abundances (<10 counts for any of the samples) were excluded from analysis. Plots were generated using ggplot2 (v3.2.1).

### 2.8 Gene set enrichment analysis and gene list generation

Published lists of upregulated genes from various recent microglia and astrocyte gene expression studies were downloaded and used for analysis (Table 1). To generate gene sets, the published gene sets were filtered to only include genes upregulated with a fold-change > 1.5 and adjusted *p* value, or false discovery rate (FDR) < 0.05. The gene lists used are available as a supplemental document. Gene Set Enrichment Analysis (GSEA) was run on full stroke IP gene lists which were ranked based on transcripts per million (TPM) using GSEA software (version 4.0.1) (Mootha et al., 2003; Subramanian et al., 2005), and GSEA was performed using 1000 permutations, weighted enrichment statistic, and Signal2Noise ranking metric against manually uploaded gene lists (Table 1).

**Table 1.**
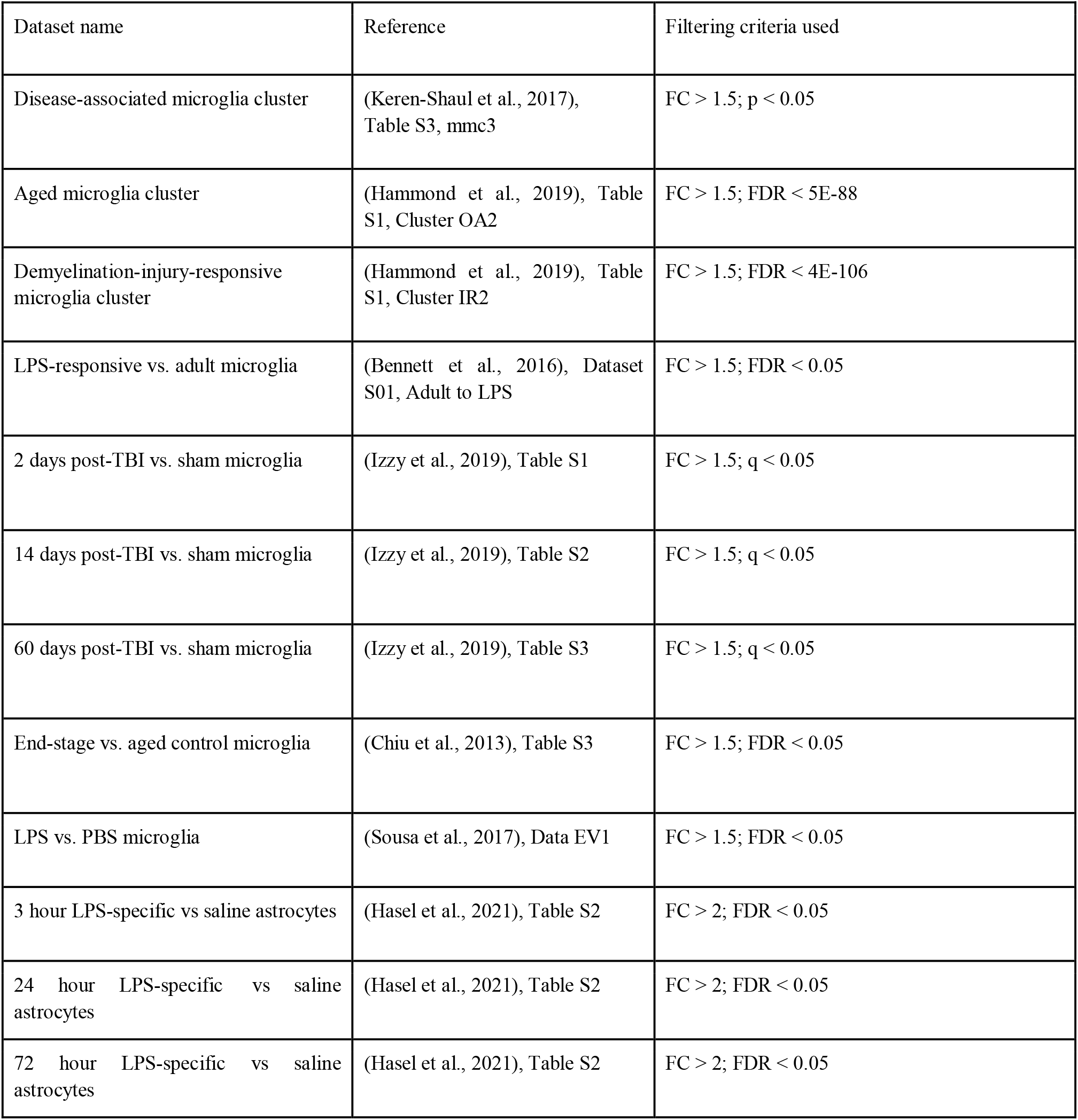
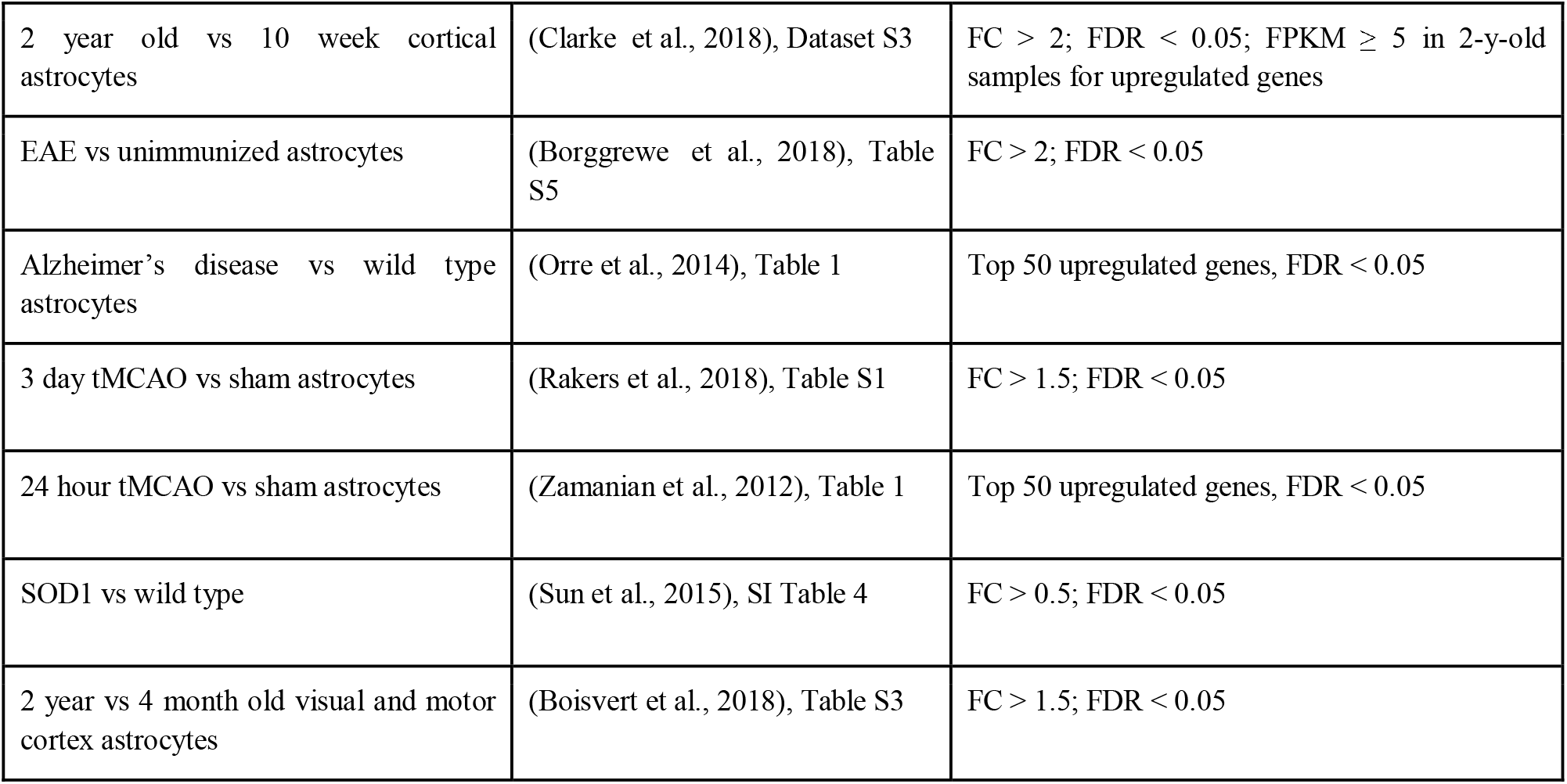
Sources and criteria for microglia and astrocyte gene sets for Gene Set Enrichment Analysis (GSEA).

### 2.9 Cell isolation for flow cytometry

Mice were deeply anesthetized 3 days after dMCAO stroke surgery, approximately 0.6 mL whole blood was obtained by cardiac puncture, then animals were transcardially perfused with cold heparinized 0.9% saline. Brains were then extracted and split into hemispheres, and cerebellum and olfactory bulbs were removed. White blood cells were obtained by red blood cell lysis and resuspension of approximately 10^6^ cells in 200 mL of HBSS. The brain hemisphere ipsilateral to the stroke was homogenized in 10 mL HBSS, filtered through a 70 μm cell strainer, and then digested in collagenase IV (1 mg/mL, Worthington, Lakewood, NJ) with DNAse in HBSS for 30 minutes at 37°C with 200 rpm shaking. Myelin was removed by addition of 5 mL of 30% Percoll (Sigma) in PBS and density-gradient centrifugation, and cells were washed in serum-supplemented DMEM. Dead cells were labeled using Live/Dead fixable aqua dead cell staining kit (ThermoFisher Scientific). Cells were fixed for 20 minutes in cold Fixation/Permeabilization buffer (BD Biosciences) then washed, and stored in fetal bovine serum containing 10% DMSO at −80°C.

### 2.10 Flow cytometry analysis

Approximately 10^6^ brain cells and 0.5 × 10^6^ blood cells were used for flow cytometry analysis. Fc receptors were blocked with 100 mL solution containing anti-mouse CD16/CD32 (1:100, BD Pharmingen, 553141) for 30 minutes at 4°C. Cells were washed and permeabilized for intracellular staining in Perm/Wash Buffer (BD Biosciences) and antibody staining was performed in 100 uL Perm/Wash buffer for 30 minutes at 4°C. The following antibodies were used for surface receptor labeling: CD45 (1:200, BioLegend, 103112), CD11b (1:200, BioLegend, 101216), HA (1:200, BioLegend, 901517), Ly6G (1:200, BioLegend, 127623), and Cx3cr1 (1:200, BioLegend, 149009). Cells were washed in buffer and analyzed with an LSR II cytometer (BD Biosciences). Gating and analysis of cell populations was carried out using FlowJo software (Tree Star Inc.).

### 2.11 Statistical analysis

Quantified immunohistochemistry data was analyzed using a two-way ANOVA with Tukey’s multiple comparisons test for post-hoc analysis in GraphPad Prism 8 software. All data is presented as mean ± SEM unless indicated. Experiments were designed using power analyses to determine sample sizes based on expected variances and group differences. All animals were randomized between experimental groups and experimenters were blinded to group assignments during analysis. Spearman correlation was calculated from the mean TPM values of genes. *Z*-scores were calculated across all IP samples for other heatmap representations. TPM values were not used for statistical analysis.

## Results

### 3.1 Characterization and validation of microglia and astrocyte RiboTag mice for cell-specific isolation of RNA after stroke

To isolate the microglia or astrocyte-specific gene expression signature after stroke, we took advantage of the genetic RiboTag technique. This allows for immunoprecipitation of ribosomes from a cell type of interest, so that the actively translating mRNA transcriptome can be isolated based on cell-type specific promoters. The Cx3cr1 promoter used in our Microglia RiboTag mice (*Cx3cr1*^*CreER2-IRES-eYFP/+*^*;Rpl22*^*HA+/+*^) (Figure 1A) is also expressed by some peripheral immune cells, including monocytes and some T cells (Jung Steffen et al., 2000; Mionnet et al., 2010; Böttcher et al., 2015). To ensure microglial-enriched mRNA and exclude infiltrating myeloid cells at the time of stroke, we allowed for 30 days to elapse between the last tamoxifen dose and stroke or sham surgery in Microglia RiboTag mice (Figure 1B). Given that peripheral immune cell populations turn over relatively quickly (Parkhurst et al., 2013; Gu et al., 2016), a 30 day wait period ensures that no peripheral Rpl22^HA^-expressing cells remain after tamoxifen administration in Microglia RiboTag mice (Supplemental Figure 1). Microglia RiboTag mice are haploinsufficient for the fractalkine receptor, but we did not observe significant weight loss after stroke (data not shown). Additionally, loss of this receptor did not affect stroke size or microglia/macrophage activation, measured by quantification of immunostaining for the microglia/macrophage activation marker CD68 (Supplemental Figure 2). The Aldh1l1 promoter is highly specific to astrocytes (Hu et al., 2019), thus we only implemented an 8 day waiting period between the last tamoxifen dose and stroke or sham surgery in our Astrocyte RiboTag mice (*Aldh1l1*^*CreER2/+*^*;Rpl22*^*HA+/+*^), to ensure tamoxifen and its metabolites were completely degraded (Valny et al., 2016).

**Figure 1.**
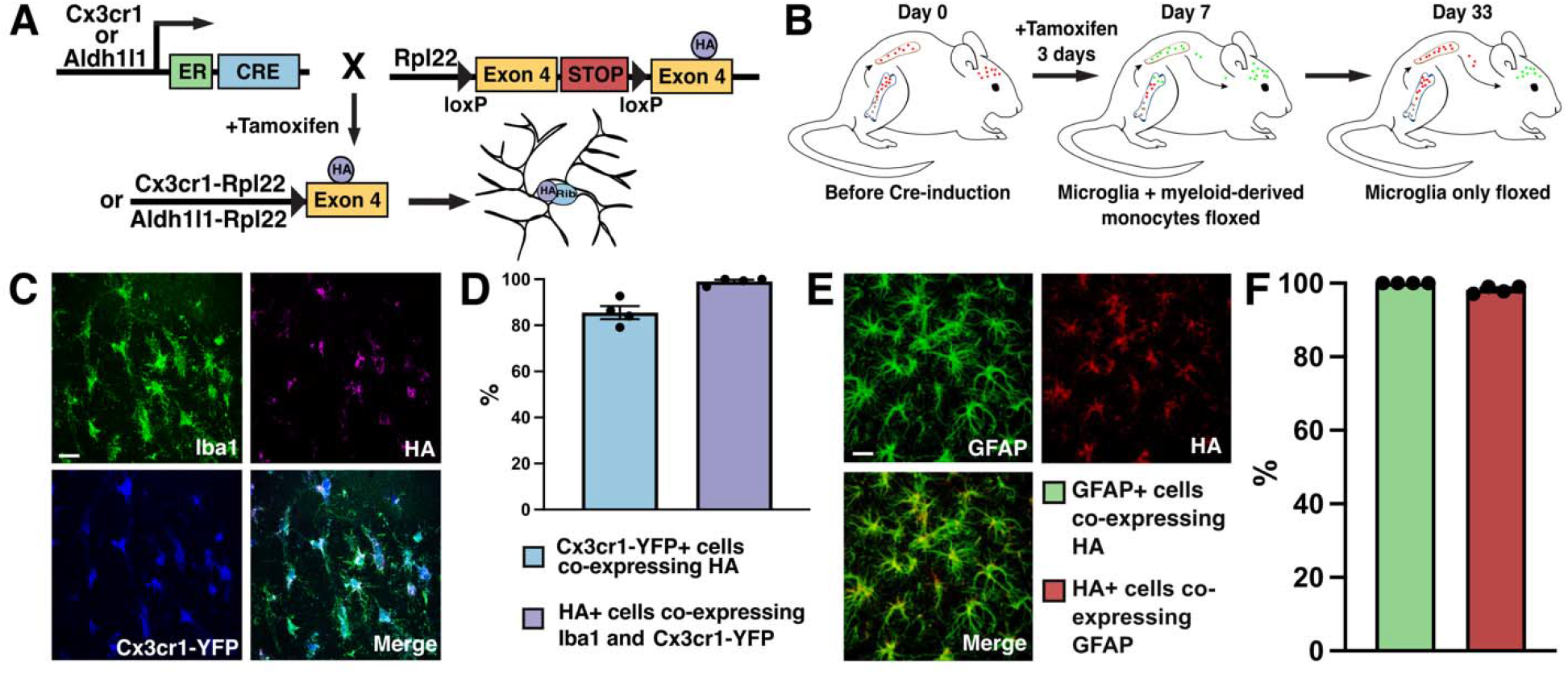
Characterization of microglia and astrocyte-RiboTag mice for isolation of microglial or astrocytic RNA after stroke. **(A)** Strategy for breeding and inducing Microglia-RiboTag (*Cx3cr1*^*CreER2-IRES-eYFP/+*^*;Rpl22*^*HA+/+*^) and Astrocyte-RiboTag (*Aldh1l1*^*/+*^*;Rpl22*^*HA+/+*^) mice. **(B)** Strategy for isolating microglia from peripheral cells in Microglia-RiboTag mice. (**C)** Representative immunofluorescence images for HA and microglia-specific markers taken in cortex of Microglia-RiboTag mice 3 days after dMCAO stroke. (**D)** Quantification of the specificity of HA by microglia and efficiency of recombination in Microglia-RiboTag mice. (99.0% +/-0.8%, *n* = 3-4). Recombination occurred in the majority of Cx3cr1+ cells quantified, and after stroke, 85.6% ± 2.8% Cx3cr1+ cells were HA+ (*n* = 4). 150 Iba1+ cells per mouse. Scale bar = 20 μm. (**E)** Representative immunofluorescence images for HA and astrocyte-specific marker taken in cortex of Astrocyte-RiboTag mice 3 days after dMCAO stroke. (**F)** Quantification of the specificity of HA by astrocytes and efficiency of recombination in Astrocyte-RiboTag mice. Almost all HA+ cells co-expressed GFAP in stroke condition (100% +/-0%, *n* = 4). Recombination occurred in the majority of GFAP+ cells quantified, after stroke, 98.2% ± 0.47% GFAP+ cells were HA+ (*n* = 4). 100 GFAP+ cells per mouse. Scale bar, 20 μm; Error Bars, SEM.

To verify faithful HA expression in our cell types of interest, we used fluorescence confocal microscopy to colocalize HA and cell-specific markers in brain sections from Microglia RiboTag or Astrocyte RiboTag mice 3 days after stroke. Almost all HA immunostaining (99%) colocalized with microglia markers Iba1 and Cx3cr1, indicating high specificity of the Rpl22 tag (Figure 1C, D). In Astrocyte RiboTag mice, 100% of HA-expressing cells co-localized with the astrocyte marker GFAP, and 98% of GFAP-expressing cells also express HA, validating that the HA tag is efficiently expressed specifically in astrocytes in this mouse line (Figure 1E, F).

### 3.2 Isolation and sequencing of microglia and astrocyte-enriched mRNA at hyperacute and acute timepoints after ischemic stroke

We immunoprecipitated microglia-enriched (microglia-IP samples) and astrocyte-enriched (astrocyte-IP samples) mRNA from the stroke core and 2-3 mm of surrounding peri-infarct cortex at 4 hours and 3 days after dMCAO stroke or sham surgery. In addition, we isolated RNA from the non-immunoprecipitated homogenate (Input samples) to assess overall gene expression and further validate the cell-specificity of our RiboTag mouse lines. For each of the two time points, we sequenced microglia-IP mRNA samples from 9-12 individual adult mice after stroke or sham surgery (Figure 2A). For the astrocyte-IP cohort, we sequenced samples from 12-20 individual adult mice after dMCAO stroke or sham surgery (Figure 2A). Half the animals used for this study were male and half were female to allow us to examine sex-specific changes in microglial and astrocyte gene expression after ischemic stroke.

**Figure 2.**
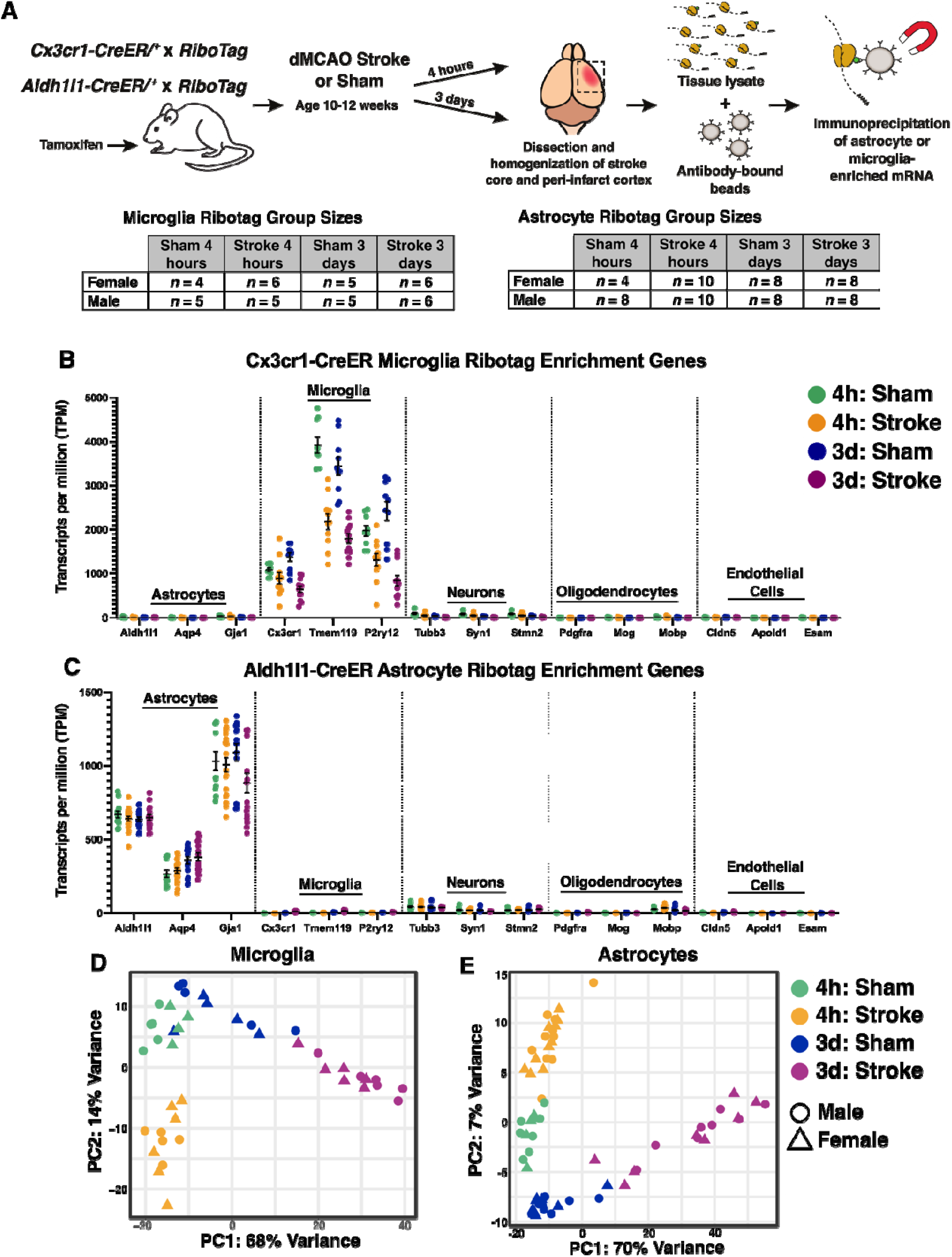
Isolation and sequencing of microglia and astrocyte-enriched mRNA at multiple time points after ischemic stroke. **(A)** Schematic depicting the experimental strategy for isolating actively translating microglial or astrocyte RNA at multiple time points after ischemic stroke, and number of biological replicates used for RNA-sequencing. All adult mice (10-12 weeks old at time of surgery) received tamoxifen prior to stroke or sham. Microglia-RiboTag mice received tamoxifen 3 days *p*.*o*. 30 days before surgery. Astrocyte-RiboTag mice received tamoxifen 5 days *p*.*o*. 8 days before surgery. Mean TPM values indicating that **(B)** microglia-IP samples are enriched in microglia-specific genes, **(C)** Astrocyte-IP samples are enriched in astrocyte-specific genes, and **(B, C)** de-enriched for cell-specific markers for other brain cell types. Bars, ± SEM. (**D)** Principal component analysis (PCA) of log-transformed RNA-seq data from 4 hour and 3 day stroke and sham microglia-IP samples (*n* = 9-12 mice per group). (**E)** Principal component analysis (PCA) of log-transformed RNA-seq data from 4 hour and 3 day stroke and sham astrocyte-IP samples (*n* = 12-20 mice per group).

To validate cell-specificity in the RNASeq data we examined cell-specific genes encompassing astrocytes, microglia, neurons, oligodendrocytes, and endothelial cells. We analyzed the expression levels of canonical brain cell markers in microglia-IP samples and astrocyte-IP samples in all four experimental groups: 4 hour & 3 day, and sham & stroke. We observed high cell-specificity in IP samples from both mouse lines (Figure 2B, C). Principal component analysis also demonstrated distinct separation of the 4-hour sham, 4-hour stroke, 3-day sham, and 3-day stroke groups in both cell types. The first two principal components accounted for 82% of the variance in gene expression among the microglia-IP samples (Figure 2D) and 77% for the astrocyte-IP samples (Figure 2E). Interestingly, for both microglia and astrocytes, the 3-day stroke and sham groups separated along a continuum in the first principal component, whereas the 4-hour stroke and sham samples clustered into discrete groups in the second principal component.

### 3.3 Differential gene expression analysis of microglial and astrocyte transcripts at two time points after stroke

To interrogate how stroke drives translatome changes, we identified differentially expressed genes (DEGs) at 4 hours and 3 days in microglia and astrocytes using the criteria of adjusted *p*-value (*p*-adj) < 0.05, and |Log2 fold change| > 1 comparing stroke versus sham. In microglia, differential expression analysis between the stroke and sham groups identified 204 unique upregulated genes and 27 downregulated genes at 4 hours, and 828 upregulated genes and 608 downregulated genes at 3 days (Figure 3A). In astrocytes, we identified 99 unique upregulated genes and 5 downregulated genes at 4 hours, and 1630 upregulated genes and 94 downregulated genes at 3 days (Figure 3B). Interestingly, the gene changes induced by stroke were fairly distinct at each timepoint in both cell types. In microglia, only 132 upregulated and 4 downregulated genes were shared between the 4 hour and 3 day timepoints. In astrocytes, the two timepoints only shared 234 upregulated genes and no downregulated genes. While the 3 day time point had a greater number of overall differentially expressed genes compared to the 4 hour time point, both microglia and astrocytes upregulated more genes than downregulated for each time point (Figure 3C-F).

**Figure 3.**
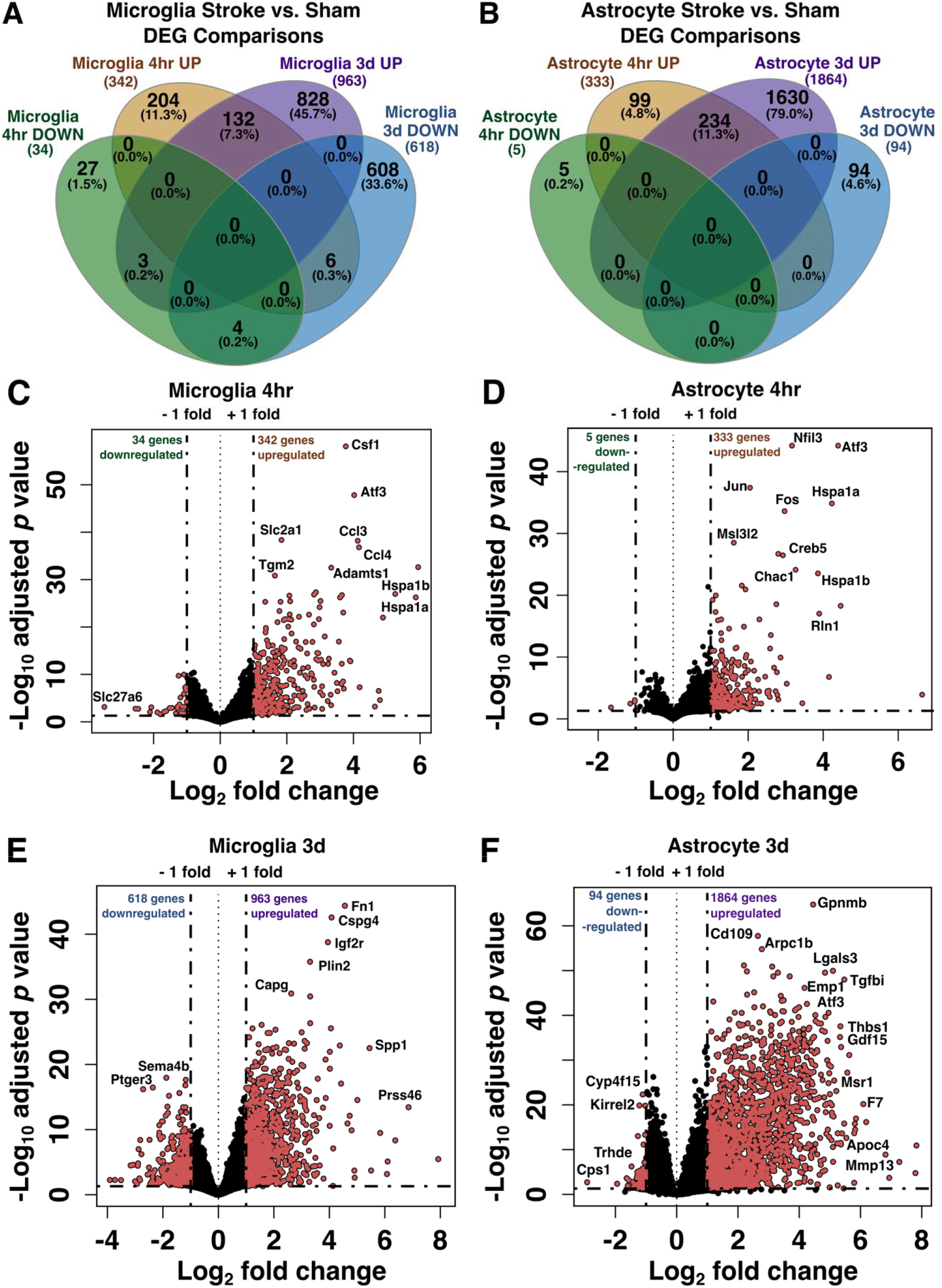
Differential gene expression analysis of microglial and astrocyte transcripts at two time points after stroke. Venn diagrams showing the number of unique and shared differentially expressed genes between stroke versus sham at 4 hours and 3 days in microglia **(A)** and astrocytes **(B)**, separated by up- and down-regulated genes (*p*-adj < 0.05; |Log_2_FC| > 1). (**C-F)** Volcano plots showing changes in microglial and astrocyte gene expression 4 hours **(C, D)** or 3 days **(E, F)** after stroke. Pink points indicate genes which passed a differential expression cutoff of *p*-adj < 0.05 and |Log_2_FC| > 1. Genes with the largest changes in expression level are labeled. Horizontal dashed line marks adjusted *p* value = 0.05; DEG, differentially expressed genes.

### 3.4 Sex differences in the microglia and astrocyte translatome after ischemic stroke

Interestingly, in both microglia and astrocytes, male and female samples did not cluster separately in the principal components analysis (Figure 2D, E), so we included all samples regardless of sex in the stroke vs. sham differential expression analysis (Figure 3). However, since stroke induces so many gene changes, it was possible that stroke-induced transcriptional changes might be masking sex differences in the principal components analysis. Therefore, to further investigate the effect of sex on translational changes in microglia-IP and astrocyte-IP samples after stroke, we directly compared female vs. male samples within each of the four experimental conditions. Surprisingly, there were still relatively few gene expression differences between female and male mice in stroke or sham conditions at both time points, and in both microglia and astrocytes (Figure 4A, B). Five sex-chromosome-linked genes, *Xist, Kdm5d, Uty, Ddx3y*, and *Eif2s3y*, were different in all conditions in both cell types. In the 4 hour sham group, only five genes in microglia and astrocytes were differentially expressed between females and males (Figure 4C, D). In the 4 hour stroke group, only three genes were differentially expressed in microglia (Figure 4E), while none were differentially expressed in astrocytes. The 3 day sham condition also exhibited few differences, with only three genes in microglia and 1 gene in astrocytes that were differentially expressed between females and males (Figure 4F, G). Sex had the greatest effect on the microglial translatome at 3 days after stroke, and interestingly, 77.8% of the genes that were different between females and males were upregulated in females (Figure 4H). These included the genes *Kcnt2, Scn8a*, and *Cacnb4*, which are known to be involved in regulation of synaptic transmission and potentiation. Conversely, in astrocytes 3 days after stroke there were few differences with only 1 differentially expressed gene in females vs. males (Figure 4I).

**Figure 4.**
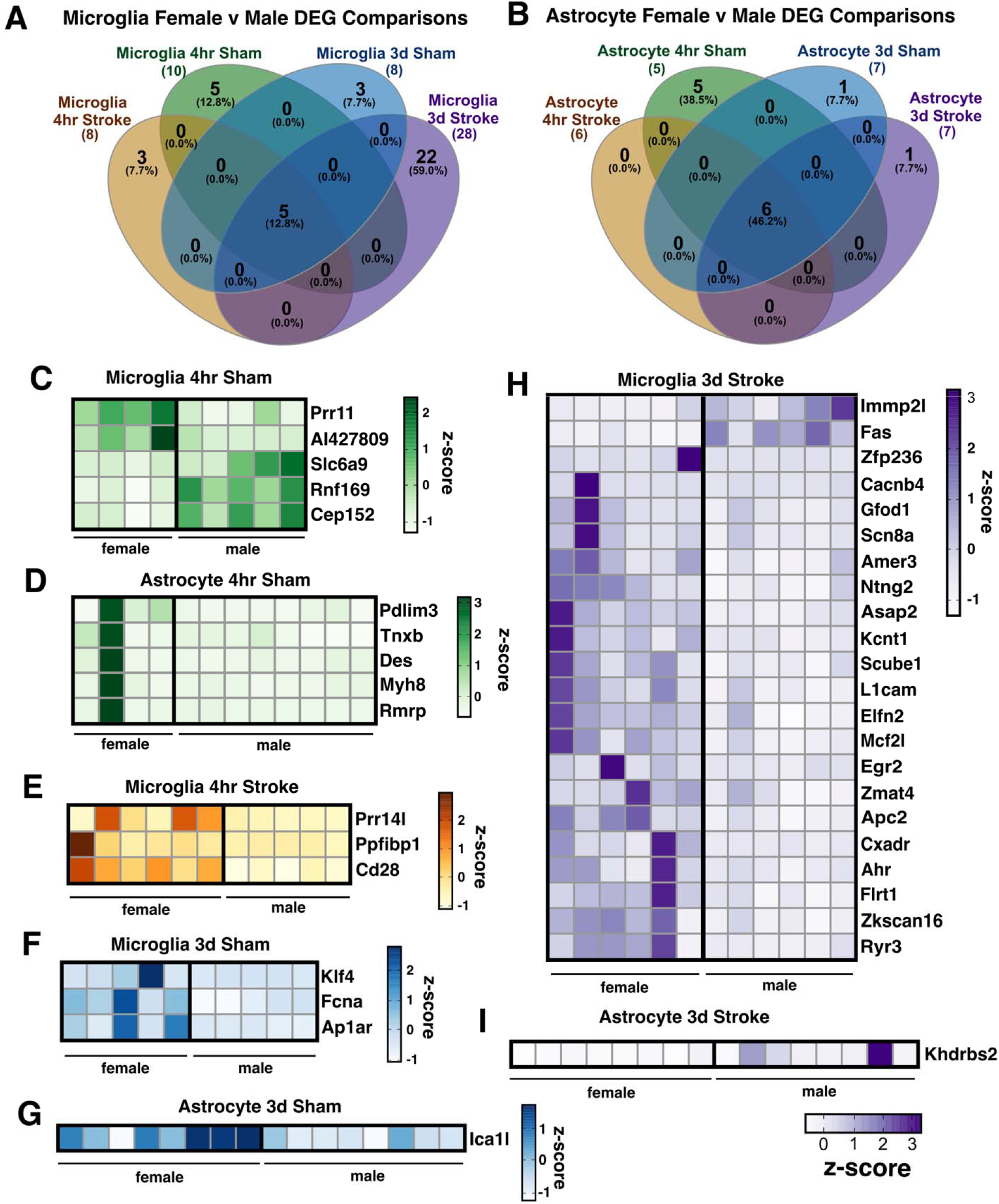
Sex differences in the microglia and astrocyte translatomes after ischemic stroke. Number of differentially expressed genes (*p*-adj < 0.05; |Log_2_FC| > 1) between males and females at 4 hours or 3 days after stroke or sham surgery in **(A)** microglia-IP samples and **(B)** astrocyte-IP samples. Heatmaps depict expression levels of significantly different genes between female and male mice unique to 4 hours after dMCAO sham **(C, D)** or stroke surgery **(E)** and 3 days after dMCAO sham **(F, G)** or stroke surgery **(H, I)**. Sex-linked genes differentially expressed by sex across all conditions were excluded from heatmaps (*Uba1y, Ddx3y, Kdm5d, Eif2s3y, Uty, Xist*). z-scores were calculated from TPM normalized gene expression values. DEG, differentially expressed genes.

### 3.5 The microglia and astrocyte translatomes reflect distinct functions during the hyperacute and acute phases of neuroinflammation after ischemic stroke

To investigate the hyperacute and acute gene expression signature unique to microglia and astrocytes after ischemic stroke, we explored the transcripts which met the criteria of being greater than 2-fold enriched in stroke versus sham conditions at 4 hours and 3 days after stroke (and significant, *p-*adj < 0.05). With the exception of microglia at 3 days, the number of significant downregulated genes was small (<100) across conditions, thus we decided to primarily focus our analysis on upregulated genes. The top upregulated genes at the hyperacute and acute timepoints suggest that microglia and astrocytes are engaged in distinct functions during the two time points. During the initiation phase of neuroinflammation after stroke, measured at 4 hours after stroke, the most highly up-regulated genes (Figure 5A) in microglia were the heat shock proteins *Hspa1a, Hspa1b*, the chemokines *Cxcl1, Cxcl2*, and macrophage inflammatory proteins *Gdf15* and *Ccl4*. These gene changes likely reflect a marked microglial response to stress and damage associated molecular patterns (DAMPs) in the ischemic region early after injury. Heat shock protein expression also likely propagates the damage signal and the inflammatory response in the injured tissue. Of the microglia genes which were most significantly regulated by stroke, *Ccl4* and *Ccl3* were expressed at very high levels, suggesting that they may be central features of the microglial stroke response at this time point.

**Figure 5.**
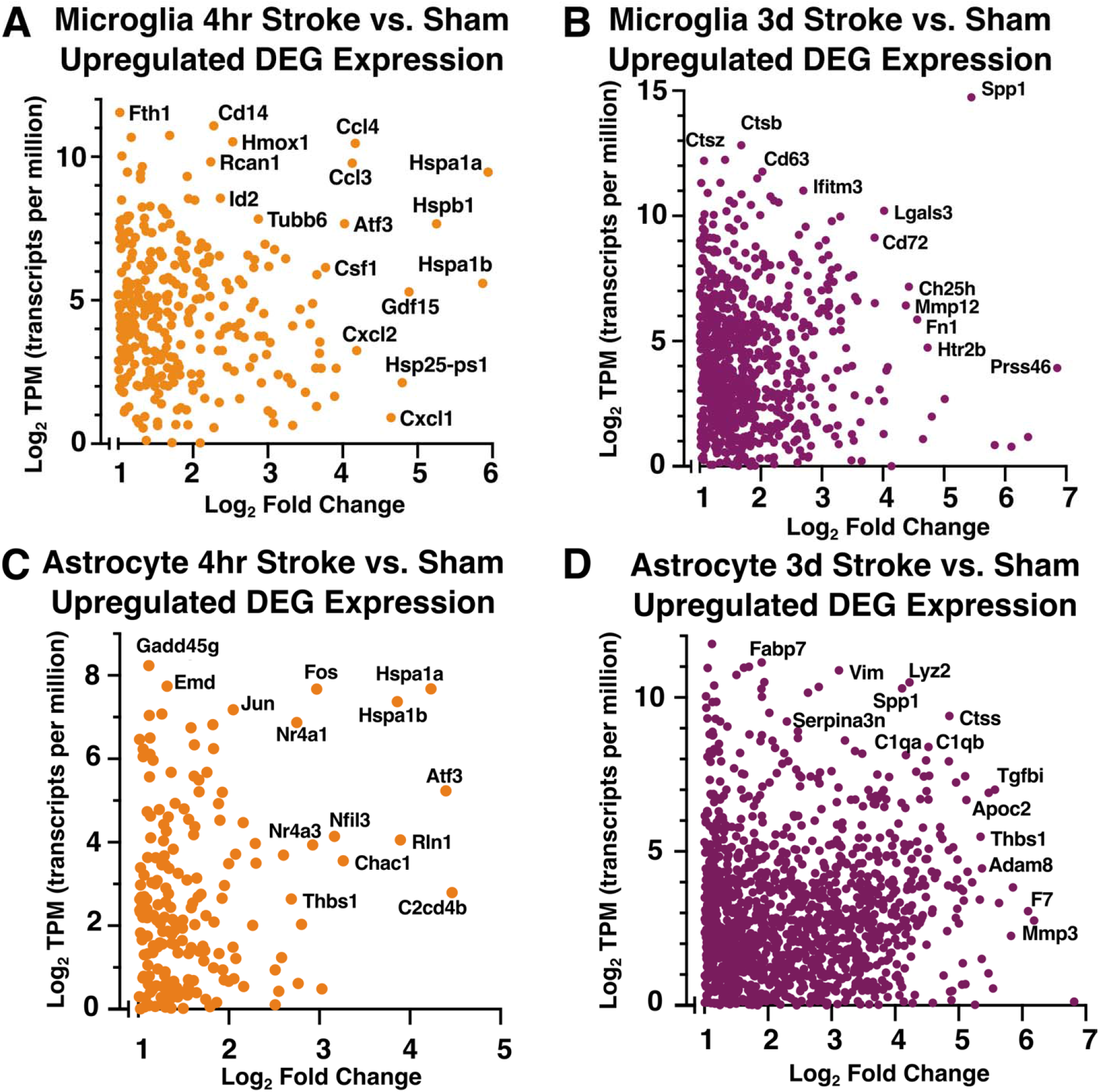
The microglia and astrocyte translatomes reflect distinct functions during th hyperacute and acute phases of neuroinflammation after ischemic stroke. **(A, B)** Scatter plots for microglia-IP **(A, B)** and astrocyte-IP **(C, D)** log_2_ mean normalized expression value (TPM) and log_2_ fold change for each significantly upregulated differentially expressed gene at 4 hours **(A, C)** and 3 days **(B, D)** after stroke compared to sham. TPM values used for this representation are the mean TPM for each gene across all samples within respective 4 hour and 3 day stroke groups. Highly expressed and/or differentially genes are labeled. Genes with TPM <1 were excluded.

During the acute phase at 3 days after stroke, *Spp1* (osteopontin 1) was highly upregulated and also had a higher expression level than any other gene at the 3 day time point (Figure 5B). *Spp1* is upregulated in degenerative disease-associated microglia (DAM) (Keren-Shaul et al., 2017), is upregulated in the brain after ischemia, and is correlated with phagocytosis (Shin et al., 2011). Also highly upregulated at this time point was *Ch25h*, the gene for cholesterol 25-hydroxylase which plays an important role in lipid metabolism, and has been linked to risk for Alzheimer’s disease (Papassotiropoulos et al., 1145059200), suggesting microglia are processing lipids engulfed during debris clearance. Other genes which were highly upregulated during the acute phase of neuroinflammation after stroke included *Lgals3* (galectin 3), which promotes microglial proliferation (Rahimian et al., 2018), and *Mmp12* (matrix metalloproteinase 12), which has been associated with worse neuronal damage after stroke (Chelluboina et al., 2015). Upregulation of phagosome-associated genes such as *Ch25h*, the macrophage scavenger receptor 1 (*Msr1*), and complement *C3* indicate that microglia regulate acute inflammation after stroke by phagocytosing dead cells and debris. Interestingly, microglia also upregulate *Cd22*, a negative regulator of phagocytosis in aged microglia (Pluvinage et al., 2019), at 3 days but not at 4 hours after stroke. It is currently unknown whether *Cd22* regulates the phagocytic capacity of microglia after ischemic cell death. At 3 days after stroke, many more genes were upregulated to a higher expression level than at 4 hours after stroke (Figure 5A, B).

In astrocytes, the top upregulated genes at the hyperacute and acute time points indicate that astrocytes initiate a stress response as early as 4 hours that continues at 3 days. At 4 hours, astrocytes highly expressed and highly upregulated (Figure 5C) *Hspa1a* and *Hspa1b*, which encode Hsp70, a chaperone protein that may be neuroprotective after stroke (Kim et al., 2020). Another highly expressed stress-response gene, *Gadd45g* (growth arrest and DNA damage inducible gamma), has increased gene and protein expression after tMCAO in mice and may predict poor outcomes after stroke in people (Simats et al., 2020). Astrocytes also upregulated a number of transcription factors at 4 hours, including *Fos, Jun, Atf3*, and *Nfil3*.

Unsurprisingly, at 3 days after stroke, two of the most highly expressed genes in astrocytes are involved in astrogliosis: *Gfap* (glial fibrillary acidic protein) and *Vim* (vimentin) (Figure 5D). Other top expressed and upregulated genes included *Il-6* (interleukin 6), *Igfbp2* (insulin-like growth factor binding protein 2), and *Mmp10* (matrix metallopeptidase 10) which are cellular senescence-associated secretory proteins in Alzheimer’s disease (Behfar et al., 2022). Another matrix metallopeptidase, *Mmp3*, is also upregulated and has been associated with increased blood brain barrier damage and greater neuroinflammatory response (Yang and Rosenberg, 2015). *Ctsd* and *Ctss* (cathepsins D and S) are also upregulated at 3 days. This could indicate astrocytes have increased extracellular matrix degradation capabilities at 3 days after stroke.

### 3.6 Comparisons to other glial states and disease models

At 3 days post-stroke, we observed a downregulation of microglial signature gene *P2ry12*, leading us to ask if microglia maintain their homeostatic gene expression signature after cerebral ischemia. We examined expression levels of select microglia signature genes and monocyte/macrophage signature genes compiled from previous studies (Hickman et al., 2013; Butovsky et al., 2014). Many homeostatic microglial signature genes, such as *Tmem119, P2ry12, Sall1*, and *Gpr34* were downregulated at both 4 hours and 3 days after ischemic stroke (Figure 6A). Downregulation of these signature genes was most pronounced 3 days after stroke, and expression levels of additional microglial signature genes, such as *Cx3cr1, Hexb*, and *Slc2a5*, were reduced at this time point. Consistent with previous reports, we observed that downregulation of the homeostatic microglial gene expression signature corresponded to upregulation of characteristic macrophage genes (Buttgereit et al., 2016; Bohlen et al., 2017). At 3 days after stroke, microglia strongly upregulated genes associated with macrophage activation/identity including *Fn1, Cd5l*, and *Saa3* (Figure 6B) (Hickman et al., 2013). Our data indicate that as early as 4 hours after stroke, microglia lose their homeostatic gene expression identity and shift towards a macrophage-like phenotype.

**Figure 6.**
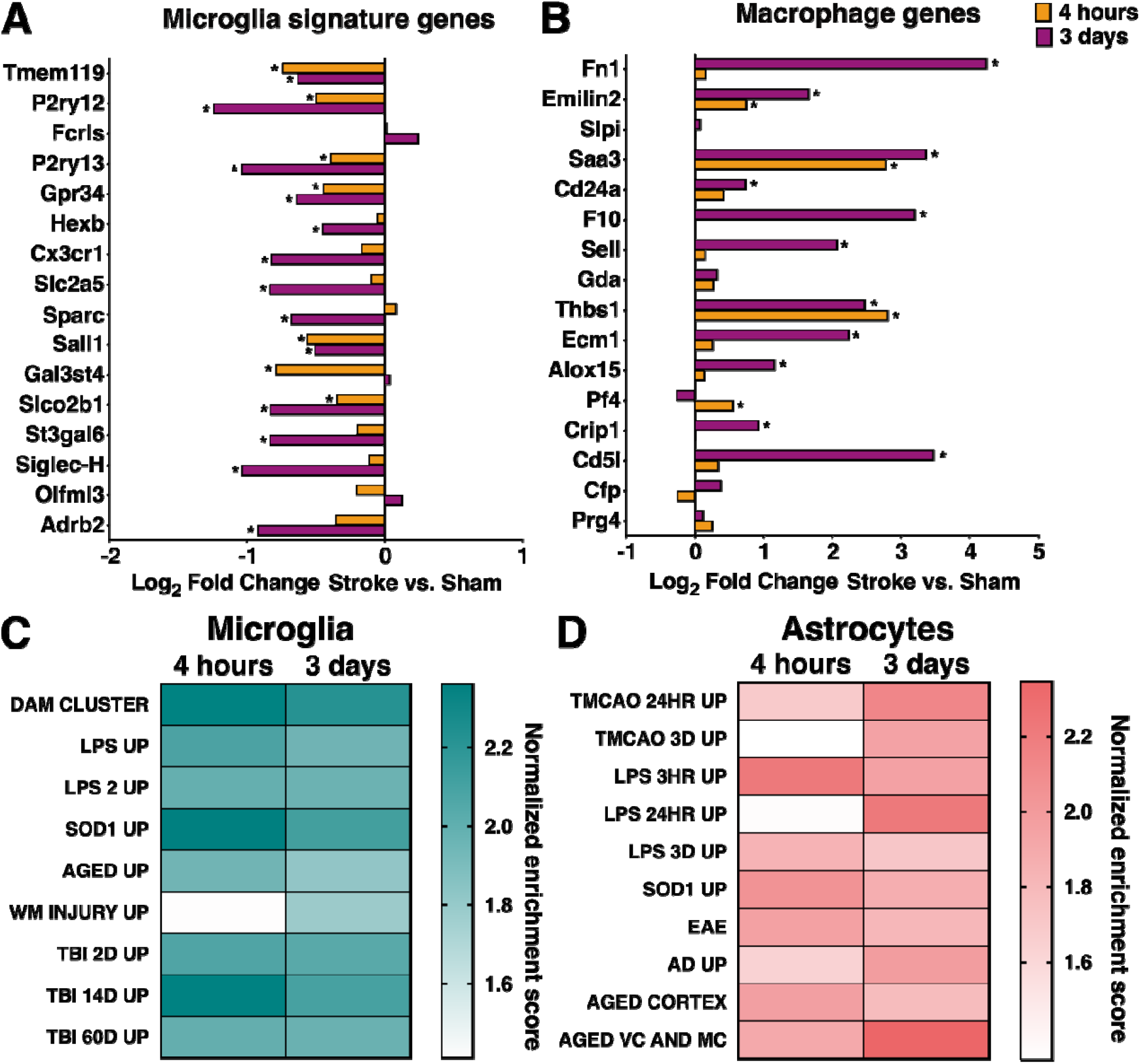
Comparisons to other glial states and disease models. Log_2_ fold change in microglial gene expression in stroke versus sham samples at 4 hours and 3 days after dMCAO stroke for select microglial signature genes **(A)** and select macrophage marker genes **(B)**. Gene set enrichment analysis results comparing microglial **(C)** and astrocyte **(D)** gene expression at 4 hours and 3 days after stroke to published datasets of microglia and astrocyte reactivity in various disease and injury states. Darker color indicates higher normalized enrichment score for each comparison. All datasets were significantly enriched in respective post-stroke glial transcript lists from both timepoints, except white matter injury (WM Injury Up) and LPS after 24 hours (LPS 24hr Up), using an FDR <25%. Table 1 lists references for gene set sources.

The downregulation of signature microglial homeostatic genes is a hallmark of disease-associated microglia (DAM). The DAM phenotype is observed in mouse models of Alzheimer’s disease (AD), tauopathy, ALS, MS, and aging (Keren-Shaul et al., 2017; Krasemann et al., 2017; Leyns et al., 2017; Olah et al., 2018; Jordão et al., 2019; Sobue et al., 2021). Based on the observation that signature microglial genes were downregulated at 4 hours and 3 days, we suspected that the microglial gene expression signature after stroke shares many similarities with microglial reactivity in other disease and injury models. To identify specific features of the microglial stroke response which might be shared by other disease/injury states, we compared our datasets to lists of genes from published microglia gene expression datasets reflecting different inflammatory states (Table 1). These included microglial gene upregulation in response to LPS, aging, degenerative disease, SOD1^G93A^ mutation, white matter injury, and traumatic brain injury (Chiu et al., 2013; Bennett et al., 2016; Keren-Shaul et al., 2017; Sousa et al., 2017; Hammond et al., 2019; Izzy et al., 2019). These datasets were selected because they represent mouse microglia gene signatures in both acute injury and degenerative disease states, and statistical information about their gene lists was publicly available. All gene sets, except white matter injury at 4 hours, were significantly enriched in stroke-upregulated genes sets from both timepoints (Figure 6C). Samples from the 4 hour stroke group had greater enrichment in genes from the LPS, neurodegeneration, and traumatic brain injury sets. This was striking, since LPS and neurodegeneration are very different inflammatory states compared to an acute sterile tissue injury like ischemic stroke. We identified several genes, including *Spp1, Tlr2, Ccl3, H2-K1*, and *Ccl4*, which are upregulated in almost all of the inflammatory models analyzed and may comprise a core microglial inflammatory gene expression signature.

Similarly, we explored how generalizable the astrocytic stroke response was to other disease/injury states, choosing astrocyte-specific datasets for comparison from a variety of inflammatory contexts (Table 1). Diseased astrocyte gene sets were from tMCAO, LPS, SOD1^G37R^ mutation, EAE, Alzheimer’s disease, and aging (Zamanian et al., 2012; Orre et al., 2014; Sun et al., 2015; Boisvert et al., 2018; Clarke et al., 2018; Rakers et al., 2019; Borggrewe et al., 2021; Hasel et al., 2021). As with microglia, almost all diseased astrocyte gene sets were enriched in our astrocyte datasets from both 4 hours and 3 days after stroke (Figure 6D). The only exception was astrocytes 24 hours after LPS. Published gene sets from LPS 3 hours and 3 days, SOD1, EAE, and aged cortex were slightly more enriched in our 4 hour upregulated astrocyte genes versus 3 days. Highly expressed shared genes included *Cebpd* and *Junb* (LPS 3 hours, SOD1, and EAE), *Cebpb* and *Gadd45g* (LPS 3 hours and SOD1), and *Fos* (LPS 3 hours and EAE). Interestingly, most of these genes are transcription factors known to induce the expression of pro-inflammatory genes. *Cebpb* and *Cebpd* promote the expression of IL-6, TNFα, and IL-1β (Akira et al., 1990; Shirakawa et al., 1993; Pope et al., 1994), and have been characterized as cortical astrocyte immediate-early genes after LPS (Cardinaux et al., 2000). The remaining gene sets (transient MCAO 24 hours, transient MCAO 3 days, LPS 24 hours, Alzheimer’s disease, and aged motor and visual cortex) were more enriched in our 3 day stroke-upregulated astrocyte gene set. Shared genes between these datasets include known reactive astrocyte markers (*Vim, Gfap, Serpina3n*), and proliferation associated (*Efemp2, Spp1*), extracellular matrix remodeling (*Timp1, Ecm1*) anti-inflammatory (*Gpnmb, Tmem176a, Tmem176b)*, and pro-inflammatory genes (*Tnfrsf12a, Ctss*). Overall, the selected microglial gene sets were more enriched in the microglia 4 hour versus 3 day post-stroke data, while the astrocyte gene sets were evenly distributed between the two astrocyte timepoints.

### 3.7 Enriched microglial and astrocyte processes and pathways at 4 hours and 3 days after stroke

Gene ontology analysis revealed that top biological processes involving the genes significantly upregulated at 4 hours after stroke in microglia included inflammatory response, response and regulation of IL-1, and transcription regulation (Figure 7A). Furthermore, strong pro-inflammatory pathways such as TNF signaling, NF-αB signaling, and Toll-like receptor signaling were highly upregulated (Figure 7B). In astrocytes at 4 hours, half of the top biological processes were shared with the top processes in microglia: inflammatory response, immune response, response to lipopolysaccharide, positive regulation of tumor necrosis factor production, and positive regulation of interleukin-1 beta production (Figure 7C). The remaining top processes were also related to a pro-inflammatory immune response. Astrocytes also shared many KEGG pathways in common with microglia at 4 hours such as TNF, NF-kappa B, MAPK, and IL-17 signaling (Figure 7D).

**Figure 7.**
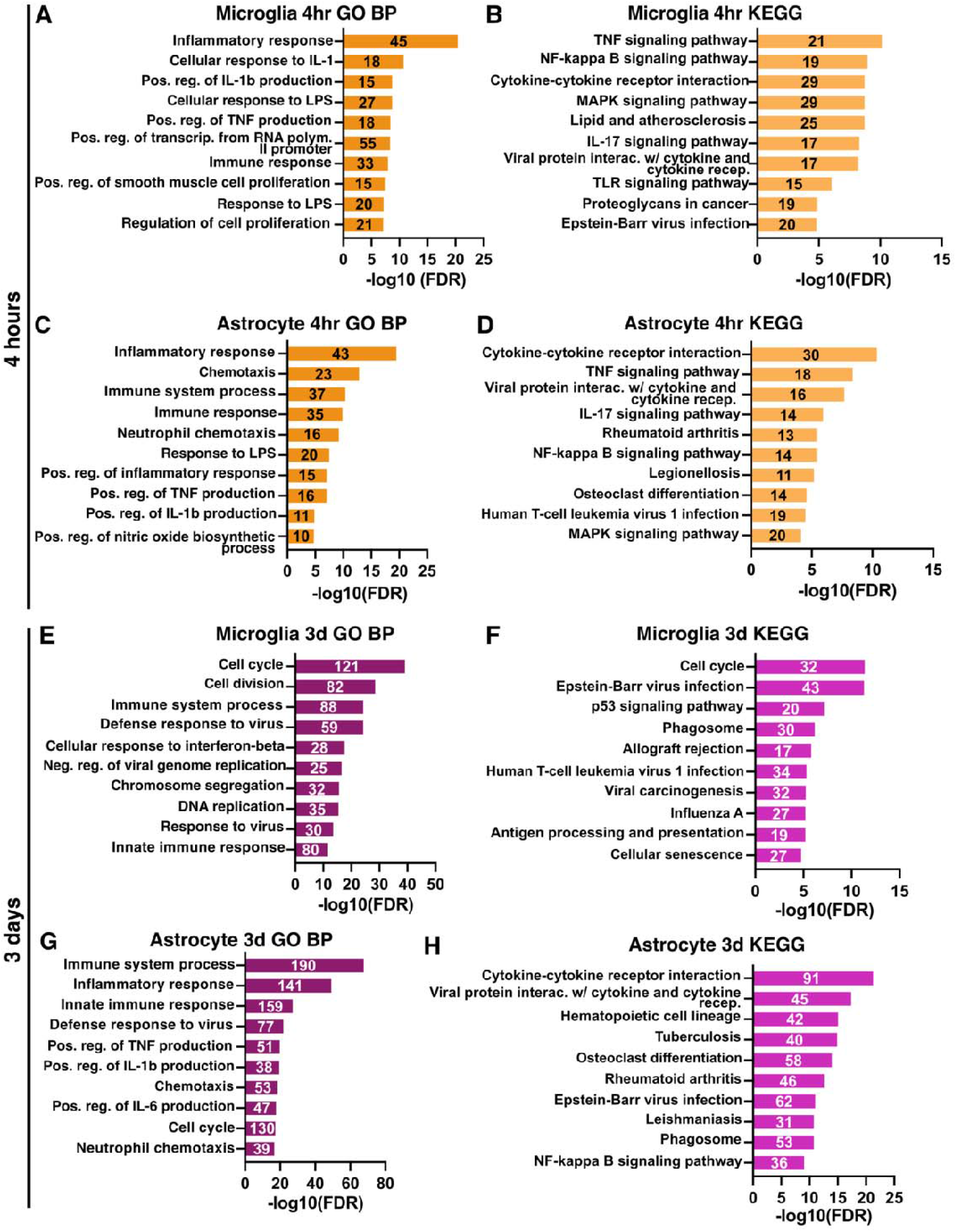
Enriched microglial and astrocyte processes and pathways at 4 hours and 3 day after stroke. Gene ontology analysis results for the top significantly enriched Biological Processes (left) and KEGG pathways (right) for microglia and astrocyte genes upregulated at 4 hours **(A-D)** and 3 days **(E-G)** after stroke. GO terms are labeled on the *y*-axis, and the number of genes from each list found in the datasets are indicated on the respective bar in the plots.

At 3 days after stroke, different pathways were identified by gene ontology analysis in both cell types. In microglia, the top biological processes and pathways reflected a functional shift and were related to cell cycling and division (Figure 7E). Immune pathways and processes also remained upregulated, including virus response genes, and phagosome-related genes (Figure 7F). In contrast, astrocytes retain a highly inflammatory profile at 3 days with most top biological processes relating to inflammation and immunity (Figure 7G). Some of the top enriched KEGG pathways were related to cytokine and cytokine receptor interactions (Figure 7H), which were also enriched at 4 hours in both cell types. However, the persistence of these pathways at 3 days could indicate that astrocytes rely more on cell-to-cell interactions at 3 days than microglia.

### 3.8 Microglia initiate an early pro-inflammatory response through TNF signaling that is propagated by astrocytes at 4 hours after stroke

At 4 hours after stroke, microglia and astrocytes shared a similar inflammatory profile, as indicated by high overlap of enriched KEGG and GO Biological Processes. This led us to interrogate how unique and shared genes in microglia and astrocytes at 4 hours contribute to the acute immune response. Using the list of genes upregulated by stroke at 4 hours in microglia (Figure 3A) and astrocytes (Figure 3B), we created a list of unique microglia genes (242 genes), unique astrocyte genes (233 genes), and shared genes by both cell types (100 genes). We performed KEGG pathway analysis on these three gene lists to obtain unique microglia (Figure 8A), unique astrocyte (Figure 8B), and shared (Figure 8C) KEGG pathways at 4 hours after stroke. Signaling pathways such as apoptosis and AGE-RAGE were uniquely enriched in the top microglia KEGG pathways. In astrocytes, top unique pathways included complement cascade, cell adhesion, and leukocyte migration. The common shared pathways reflected many of the similarities observed using all upregulated genes at 4 hours for each cell type (Figure 7B, D): TNF, IL-17, MAPK, and NF-κB signaling pathways.

**Figure 8.**
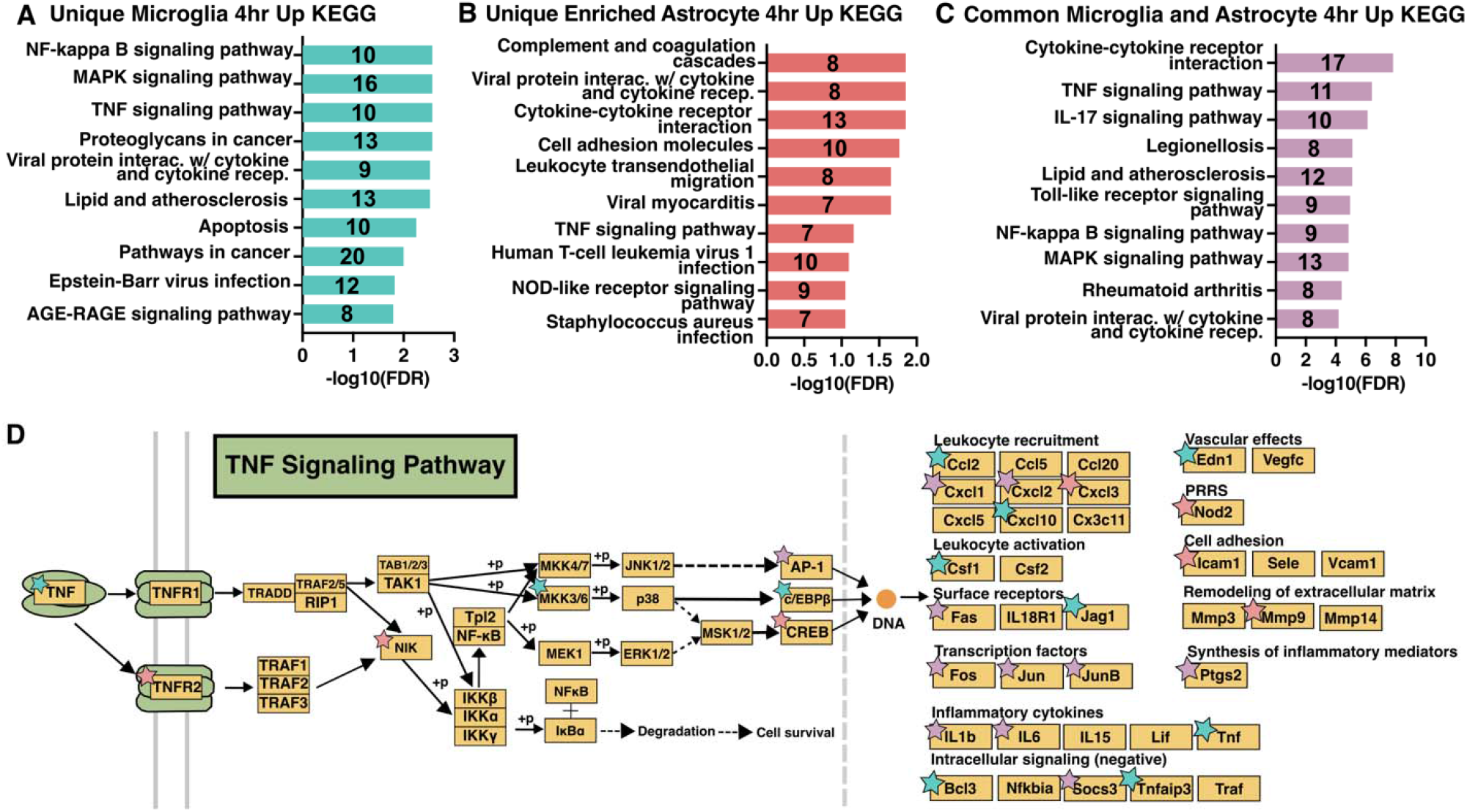
Microglia initiate an early pro-inflammatory response through TNF signaling that is propagated by astrocytes at 4 hours after stroke. KEGG Pathways analysis results for genes upregulated 4 hours after stroke and unique to microglia **(A)** or astrocytes **(B)**, or common to both **(C)**. GO terms are labeled on the *y*-axis, and the number of genes from each list found in the datasets are indicated on the respective bar in the plots. **(D)** KEGG TNF signaling pathway schematic, adapted from Kanehisa Laboratories (Kanehisa and Goto 2000), with stars denoting unique enriched upregulated genes in microglia (blue), astrocytes (pink), and shared common (purple) at 4 hours after stroke.

One advantage of obtaining microglia and astrocyte-enriched translatomes at the same time points is the ability to examine how each cell contributes to pathways. After stroke, the TNF signaling pathway in both microglia and astrocytes has been found to disrupt blood brain barrier function and worsen outcomes (Chen et al., 2019; Li et al., 2022). Because it was enriched in all three gene lists and is important in pro-inflammatory signaling, we chose the KEGG TNF signaling pathway (Kanehisa and Goto, 2000) to highlight the ability of our data to identify unique microglial, astrocyte, and shared gene contributions. TNF itself was uniquely upregulated by microglia at 4 hours after stroke (Figure 8D). All three downstream transcription factors (TFs) in the TNF pathway (AP-1, CREB, and C/EBPβ) were upregulated by either or both cell types. These then modulate downstream target genes with distinct functions (right side of Figure 8D). For each functional category there were genes upregulated in either or both cell types, revealing differential regulation within astrocytes and microglia. Notably, leukocyte activation and vascular effects were uniquely upregulated by microglia, while pattern recognition receptors, cell adhesion, and remodeling of the extracellular matrix were uniquely upregulated by astrocytes.

### 3.9 Microglia and astrocytes differentially regulate transcription at the hyperacute and acute period after stroke

To understand how astrocytes and microglia initiate and propagate inflammation, we used the transcription factors-target database TRRUST (Han et al., 2018) to identify transcription factors and their targets across time points. In the 4 hour transcription factor-4 hour target relationships (Figure 9A, B), microglia had greater numbers of upregulated transcription factors (19), targets (36), and number of interactions (53) compared to astrocytes (11 transcription factors, 22 targets, 35 interactions). However, *Jun* (jun proto-oncogene, AP-1 transcription factor subunit) and *Egr1* (early growth response 1), were upregulated in both microglia and astrocytes, and together contributed to 37.7% and 45.7% respectively of the total number of interactions at 4 hours.

**Figure 9.**
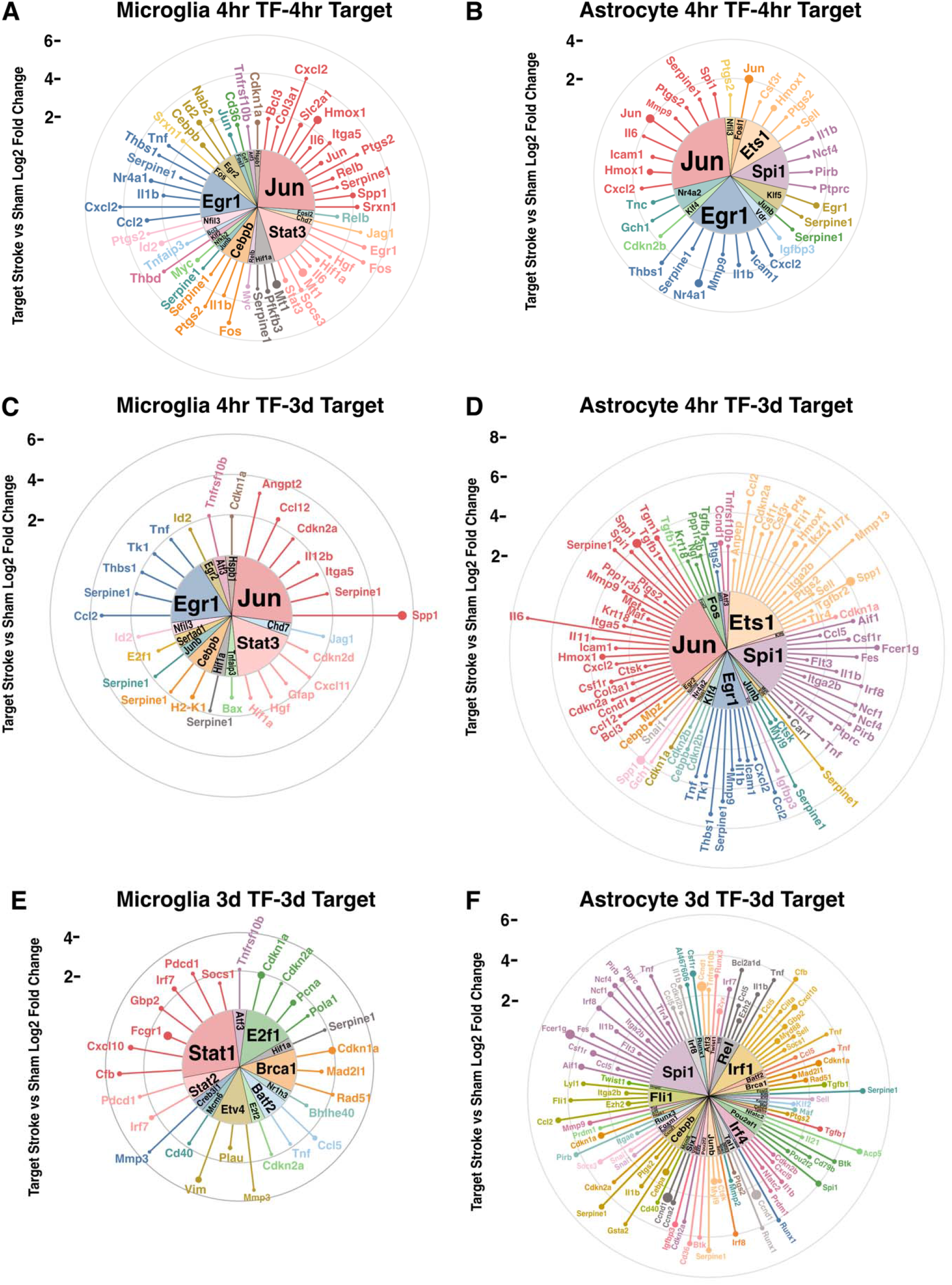
Microglia and astrocytes differentially regulate transcription at the hyperacute and acute period after stroke. Circular lollipop plots illustrating upregulated transcription factors and their upregulated targets at 4 hours and 3 days. Upregulated transcription factor-target relationships were analyzed for 4 hour transcription factors and 4 hour targets (**A, B**), 4 hour transcription factors and 3 day targets (**C, D**), and 3 day transcription factors and 3 day targets (**E, F**). Inner wedges represent the upregulated transcription factors, while each line represents its upregulated target. Length of each line indicates the log2 fold change of the target in stroke versus sham, while the size of the circle at the end of line indicates the TPM of the target.

We were also interested in how gene regulation in the hyperacute 4 hour period could affect gene expression at the acute 3 day period (Figure 9C, D). Here we observed a shift to astrocytes expressing more transcription factors (18), targets (61), and interactions (90) compared to microglia (13 transcription factors, 23 targets, and 29 interactions). The targets of *Jun* and *Egr1* remain upregulated at 3 days, as demonstrated by *Jun* and *Egr1* contributing 41.4% and 36.7% of all transcription factor-target interactions in microglia and astrocytes respectively. In astrocytes, *Ets1* (ETS proto-oncogene 1, transcription factor) and *Spi1* (Spi1 proto-oncogene) also emerged as prominent drivers of upregulated targets.

Finally, we looked to see how upregulated transcription factors at 3 days relate to targets at 3 days (Figure 9E, F). The pattern of astrocytes expanding their transcription factor-target interactions continued with 41 transcription factors, 67 targets, and 99 interactions compared to the 12 transcription factors, 22 targets, and 28 interactions in microglia. *Stat1* (signal transducer and activator of transcription 1) emerged as the dominant TF in microglia, while *Spi1* continued as a top transcription factor in astrocytes. Interestingly, in all relationships except 3 day astrocyte transcription factor-target, the number of upregulated transcription factors remained below 20, with the number of targets largely contributing to changes in total interactions. However, in astrocytes at 3 days, the number of transcription factors greatly increases, with the average number of targets per transcription factor decreasing. Overall, we infer that microglia exhibit transcriptional regulation in response to stroke to a greater extent in the hyperacute period, while astrocytes use it in the acute period.

## 4 Discussion

We present a comprehensive molecular signature of hyperacute (4 hours) and acute (3 days) transcriptional responses to stroke in microglia and astrocytes, two critical brain resident glial cell types for post-stroke neuroinflammation. To our knowledge, this is the first comprehensive characterization of microglial and astrocyte gene expression changes during the hyperacute period (4 hours) after stroke. To improve the value of the data, we constructed a user-friendly website (https://buckwalterlab.shinyapps.io/AstrocyteMicrogliaRiboTag/) where researchers can easily look up gene changes.

We report that at 4 hours after stroke, microglia and astrocytes upregulate genes characteristic of a distinctly pro-inflammatory response, including chemokines, heat shock proteins, transcription factors, and other signaling molecules which amplify inflammation. These results suggested that early after stroke, microglia are strongly proinflammatory and are particularly involved in activities such as propagation of inflammation and regulation of transcription. During the acute phase of neuroinflammation, microglia maintain a proinflammatory gene expression signature, but are strongly proliferative and entering a phagocytic state. In contrast, astrocytes appear to be similarly proinflammatory at 4 hours, but rather than shift to a predominantly proliferative state like microglia, they remain in a proinflammatory state at 3 days. In the top GO Biological Processes, 4 hour and 3 day astrocytes share 60% of processes, all related to inflammation. By comparison, 4 hour and 3 day microglia did not share any top GO Biological Processes, indicating how the microglial profile drastically shifts while the astrocyte one maintains and strengthens an inflammatory signature.

Cumulatively, the gene expression changes we observed at 4 hours after stroke suggest that microglia and astrocytes acutely respond to cerebral ischemia by turning on a distinct inflammatory gene expression program. Some of the most highly expressed and significantly upregulated genes in both cell types at this early time point were heat shock and stress response proteins, which may protect microglia and astrocytes by functioning as molecular chaperones during ischemic stress (van der Weerd et al., 2005; Sun et al., 2006; Yenari et al., 2008). The microglia specific profile at 4 hours was composed of pro-inflammatory cytokines and chemokines, including *TNF*α, *Ccl2, Ccl3, Ccl4*, and *Cxcl2*. Key functions of these molecules are positive regulation of the inflammatory response and recruitment of monocytes and neutrophils (Campbell et al., 2007; Mirabelli-Badenier et al., 2011; Wu et al., 2015). Therefore, it appears that one of the main functions of microglia early after stroke is to initiate the recruitment of peripheral immune cells into the damaged brain.

We report that there is a more similar canonical pro-inflammatory response in microglia during the hyperacute stroke response, rather than at 3 days. Ccl3 and Ccl4 were recently identified as markers of aged and neurodegeneration-associated microglia (Hammond et al. 2019; Kang et al. 2018), but the function of these chemokines in aging and neurodegeneration is unknown, and it is unclear if Ccl3 and/or Ccl4 signaling might have a unique role in the neuroinflammatory response after stroke. Our results reveal distinct microglial translational footprints at 4 hours versus 3 days after ischemic stroke, but our analysis also highlights that many of the core genes upregulated at both time points after stroke are also upregulated by microglia in various other disease and inflammatory models.

Astrocytes also display common responses in multiple disease contexts, albeit to a lesser extent compared to microglia. *Gpnmb* was shared by the largest proportion of datasets (LPS 24 hours, transient MCAO 24 hours, transient MCAO 3 days, and aged motor and visual cortex), which spans from acute to chronic inflammatory states in the brain. *Gpnmb* has been found to attenuate neuroinflammation in vitro (Neal et al., 2018) via CD44 signaling, which is most highly expressed in our data at 3 days after stroke in astrocytes. While *Gpnmb* has largely been found to be anti-inflammatory in certain pathological conditions, there is also evidence of pro-inflammatory activity (Saade et al., 2021) in others, calling for the need for further exploration. Other shared genes, *Ctss* and *Spp1*, have both been found to disrupt the blood brain barrier after stroke, with neutralization leading to improved outcomes (Wu et al., 2021; Spitzer et al., 2022; Xie et al., 2022). Overall, the astrocyte response at both time points shares some commonalities with other disease models, but also maintains a unique stroke signature.

In addition to commonalities between disease states, microglia and astrocytes display commonalities to each other in stroke. There was a high overlap of top KEGG pathways enriched in upregulated genes in both cell types in the hyperacute period. This included the pro-inflammatory pathways IL-17, NF-κB, MAPK, TLR, and TNF signaling. The TNF signaling pathway was of particular interest given that activated microglia can induce reactive astrocytes by secreting TNF (Liddelow et al., 2017). In our data, microglia began expressing *Tnf* at 4 hours after stroke, the initiator of the TNF signaling pathway, while astrocytes expressed downstream TNF pathway transcription factors such as *Fos, Jun*, and *Creb*. Thus, the two cell types may be working in concert to execute the TNF response. Also, while microglia have been established as immediate responders to injury by rapid migration to the site of tissue (Yoon et al., 2015), astrocytes also begin a unique injury response as early as 4 hours after stroke. Pro-inflammatory factors such as TNF-α secreted from microglia may initiate the pro-inflammatory signaling pathway that astrocytes respond to through upregulation of transcription factors early on to ramp up their immune response at later time points.

Transcriptional differences were greater at 3 days after stroke in both microglia and astrocytes compared to sham animals during the acute phase (3 days post-stroke) of neuroinflammation. This is not surprising, because inflammation approaches maximum levels at 3 days in the dMCAO stroke model (Chaney et al., 2019). Specific microglial gene expression changes at 3 days after stroke indicated a more complex response, and involved many genes associated with proliferation, phagocytosis, and sustained inflammation, suggesting that microglia continue to regulate neuroinflammation throughout the acute period after stroke. Microglia are less susceptible to ischemic damage than neurons and astrocytes, but studies indicate a loss of microglia in the ischemic region during the first 2 days after stroke, followed by gradual rebound in numbers over about a week (Ritzel et al., 2015; Liu et al., 2019). The proliferative transcriptional signature in our dataset likely reflects both a response to injury as well as this repopulation response.

By 3 days after stroke, astrocytes intensify the pro-inflammatory profile initiated at 4 hours into a primarily immune-driven response. In the top 10 GO Biological Processes, 7 were related to an immune or pro-inflammatory response. Specifically, genes associated with the production of pro-inflammatory cytokines Il-1β, TNF-α, and Il-6 were upregulated. Compared to 4 hours, stroke upregulated a greater number of transcription factors and their targets at 3 days in astrocytes after stroke. Of particular interest are transcription factors *C/EBPβ, Spi1*, and *Rel*, which account for 25% of all upregulated transcription factor-target interactions and are each strong candidates as therapeutic targets. Absent or reduced *C/EBPβ* has been directly associated with smaller infarct sizes, diminished cognitive deficits, reduced inflammation after tMCAO (Kapadia et al., 2006), and increased angiogenesis after stroke (Li et al., 2020). *Spi1* has also been found to be upregulated in astrocytes 3 days after tMCAO (Rakers et al., 2019) and in bulk tissue after tMCAO (Stevens et al., 2011; Vartanian et al., 2011; Zhang et al., 2019), has been identified as a regulator of astrocytic wound responses (Burda et al., 2022), and can influence NF-κB signaling (Jin et al., 2011). Another member of the NF-κB signaling cascade, *Rel*, has been shown to play a role in hippocampus-dependent memory formation (Aleyasin et al., 2004; Levenson et al., 2004) and can confer cellular resistance to hypoxia (Qiu et al., 2001). Thus, *Rel* is a promising candidate for amelioration of post-stroke cognitive deficits. It may be that the initiation and persistence of these transcription factors are key for determining proper resolution of the immune response to stroke. Importantly, we report substantial differences between both cell types’ gene expression during hyperacute versus acute neuroinflammation after stroke. Specifically, microglia respond rapidly by upregulating transcription factors and shift away from a purely pro-inflammatory state by 3 days after stroke, whereas astrocytes remain in a pro-inflammatory state across the acute phase.

Surprisingly, we found few sex-differences in the microglial and astrocyte response. Brain-wide changes in gene expression as well as neuroprotective capacity of microglia (Villa et al., 2018) and astrocyte reactivity (Cordeau et al., 2008) have been reported by others to be dependent on sex after stroke. And there is clear data on sex-dependent differences in outcomes (Branyan and Sohrabji, 2020; Noh et al., 2023). There are also significant sex differences in adult microglial gene expression in wildtype mice (Guneykaya et al., 2018), under germ free conditions (Thion et al., 2018), after LPS stimulation (Hanamsagar et al., 2017), and in astrocyte gene expression in EAE (Tassoni et al., 2019), early development (Rurak et al., 2022), and in humans (Krawczyk et al., 2022). It may be that the gene differences we did observe in microglia may drive differences in stroke outcomes in females, or that these are less prominent in the dMCAO model because it does not model reperfusion. In addition, differences in the gene expression profiles reported by us compared to others may be due to differences in isolation techniques. It may be that microglial isolation by cell sorting skews gene expression towards a pro-inflammatory state and accounts for the higher representation of pro-inflammatory genes in other studies compared to ours (Haimon et al., 2018; Kang et al., 2018; Kronenberg et al., 2018; Rajan et al., 2019) or that sorting-based isolation may differentially affect gene expression in male versus female microglia. Data on sex differences in astrocyte gene expression is limited, but (Rakers et al., 2019) also used both sexes using a ribosomal pulldown method to isolate astrocyte RNA after stroke and no sex differences were reported. This is thus an area which requires further investigation.

Indeed, we chose the RiboTag method to avoid known gene changes from cell dissociation techniques that may confound stroke-induced changes particularly at the 4 hour time point. The 4 hour acute time point is critical because it is a clinically accessible time point for delivery of future therapies, and our data provides the first published resource describing it. The next earliest astrocyte and microglial transcriptomes after stroke were 12 hours after stroke (Ma et al., 2022; Zheng et al., 2022). Other advantages of our study are the high number of biological replicates, use of both sexes, and high sequencing depth (>12 million reads per sample). Our data provide a starting point for identifying candidate pathways and signaling cascades that microglia and astrocytes may be using in a complementary fashion. Caveats to this dataset include that it is in adult mice and does not model the reperfusion injury that is part of some strokes. Also, microglia and astrocytes were not sequenced from the same mice. Although there has been a rise in global stroke burden in young and middle-aged adults (Feigin et al., 2014), aged mice will need to be incorporated in future stroke transcriptomic studies. Interestingly, our gene set enrichment analysis demonstrated that our 3 day astrocyte DEGs were enriched in a healthy, aged astrocyte population (Boisvert et al., 2018).

In summary, our results show that ischemic stroke induces substantial gene expression changes in microglia and astrocytes very early after injury that continue thereafter. Importantly, we found differences in the response profiles at each of these time points, supporting the concept that microglia and astrocytes play distinct roles in both initiation and regulation of neuroinflammation after stroke. Our data provide a useful resource for future studies investigating the functions of microglia and astrocytes in the response to ischemic stroke and may facilitate exploration of new targets for stroke treatments. For example, our analysis of the TNF pathway demonstrates the utility of this dataset in dissecting cell-specific responses to stroke. By examining gene changes in our user-friendly website (https://buckwalterlab.shinyapps.io/AstrocyteMicrogliaRiboTag/), we hope that stroke researchers will be able to generate hypotheses to speed our understanding of the neuroinflammatory response to stroke and develop therapies. Future work will also investigate how particular targets identified by our analysis affect stroke outcomes. Additionally, these experiments point to the usefulness of the RiboTag method for studying the time course of injury responses in the central nervous system.

## Acknowledgments

This work was funded by the National Institutes of Health R01NS067132 to MSB and T32MH020016 to VGH, American Heart Association/Allen Frontiers Group Brain Health Award 19PABHI34580007 to MSB, Leducq Foundation Transatlantic Network of Excellence 19CVD01 to MSB, and Stanford Graduate Fellowship and Horatio Alger Dennis Washington Leadership Graduate Scholarship to VGH. We would like to acknowledge Karen Bradshaw, Kristy Zera, and Elizabeth Mayne for their help with manuscript revisions, and Kanehisa Laboratories for granting copyright permission for the TNF signaling pathway.

## Notes

**Conflict of interest statement:** The authors declare no competing financial interests.

https://buckwalterlab.shinyapps.io/AstrocyteMicrogliaRiboTag/

### Competing Interest Statement

The authors have declared no competing interest.

### Summary of Updates

Author list has been updated

