## Supplemental Figures for "Translatome analysis reveals microglia and astrocytes to be distinct regulators of inflammation in the hyperacute and acute phases after stroke"

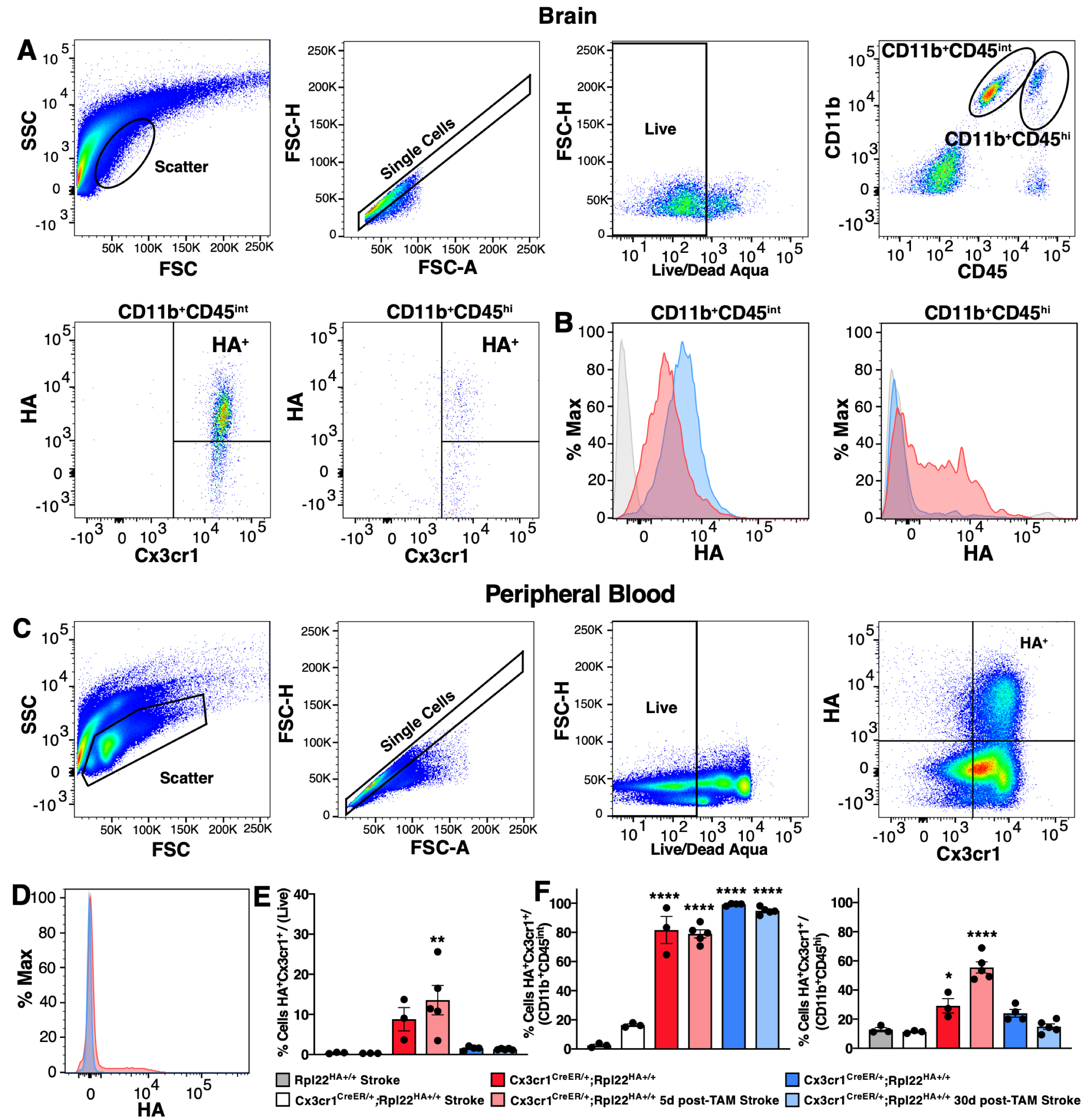

**Supplemental Figure 1. The HA tag is not expressed by peripheral immune cells 30 days after tamoxifen treatment in *Cx3cr1^CreE2R/+^;Rpl22^HA+/+^* mice. (A)** Flow cytometry gating strategy to evaluate HA expression in brain of *Cx3cr1^CreE2R/+^;Rpl22^HA+/+^* mice 5 days after tamoxifen treatment and 3 days after dMCAO stroke. **(B)** Analysis showing HA staining in CD11b^+^CD45^int^ (left) and CD11b^+^CD45^hi^ (right) cell populations isolated 3 days after stroke from the brains of control mice (grey histogram), or *Cx3cr1^CreE2R/+^;Rpl22^HA+/+^* mice 5 days (red histogram) or 30 days (blue histogram) after tamoxifen treatment. **(C)** Flow cytometry gating strategy to evaluate HA expression in peripheral blood of *Cx3cr1^CreE2R/+^;Rpl22^HA+/+^* mice 5 days after tamoxifen treatment and 3 days after dMCAO stroke. **(D)** Analysis showing HA staining in live subset of peripheral blood cells isolated 3 days after stroke from the brains of control mice (grey histogram), or *Cx3cr1^CreE2R/+^;Rpl22^HA+/+^* mice 5 days (red histogram) or 30 days (blue histogram) after tamoxifen treatment. **(E)** Percentage of live peripheral blood cells which are Cx3cr1^+^HA^+^ (*n*=3-5). One-way ANOVA with Tukey’s multiple comparisons test. Bars, mean ± SEM; ***p* < 0.01 relative to *Rpl22^HA+/+^ Stroke* condition.  **(F)** Percentage of CD11b^+^CD45^int^ (left) and CD11b^+^CD45^hi^ (right) cells which are Cx3cr1^+^HA^+^ in the brain. One-way ANOVA with Tukey’s multiple comparisons test (*n* = 3-5 mice per group). Bars, mean ± SEM; **p* < 0.05, *****p* <0.0001 relative to *Rpl22^HA+/+^ Stroke* condition.

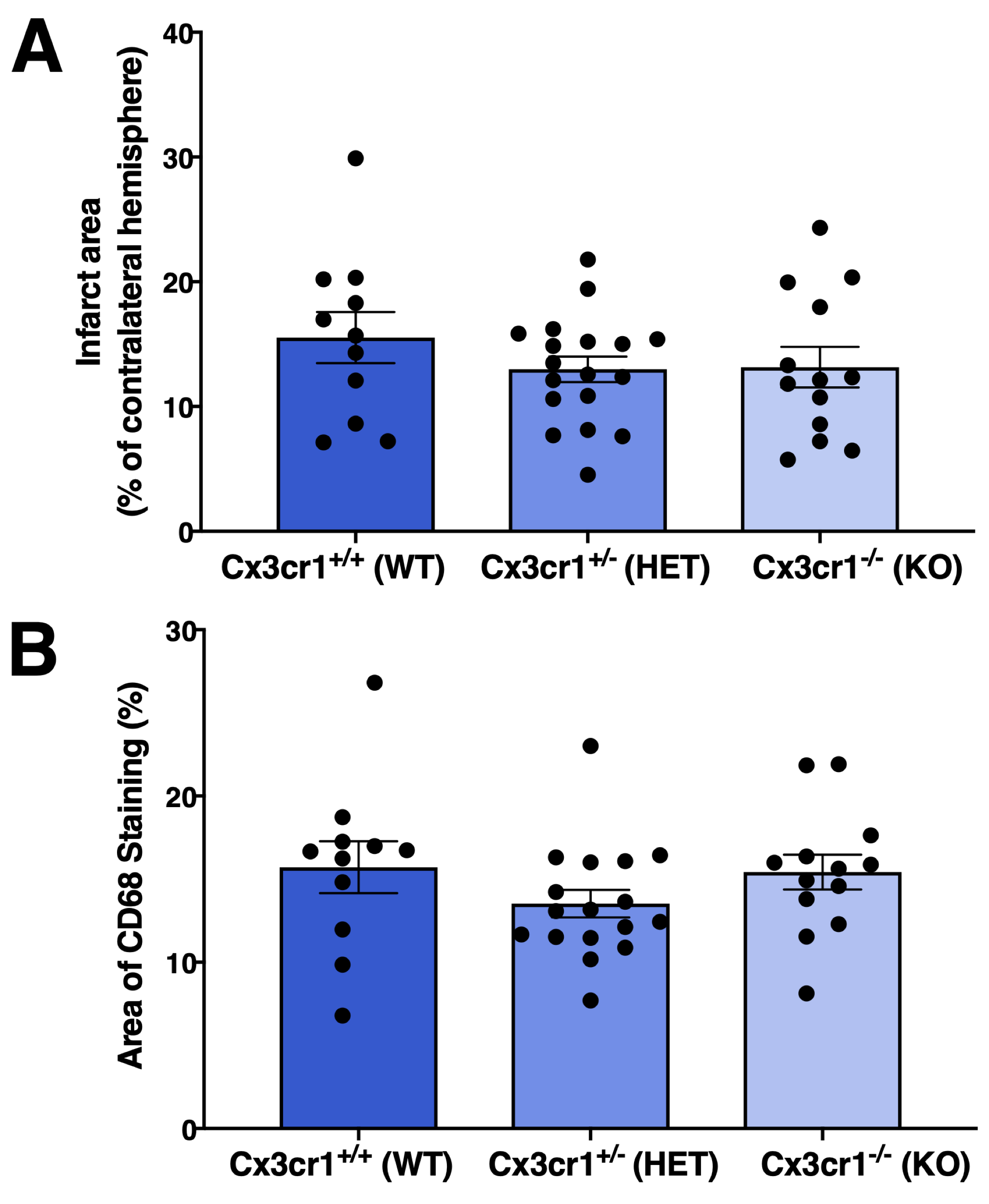

**Supplemental Figure 2. Cx3cr1 haploinsufficiency does not affect stroke size or microglia/macrophage activation in the stroke border in the dMCAO model. (A)** Quantification of infarct size (represented as percentage of contralateral hemisphere) at 3 days after stroke. **(B)** Quantification of the percent area covered by CD68 immunostaining in images taken in the stroke border. For all three measures, no significant differences were found between wildtype, heterozygous, or knockout groups (*n* = 11-18).
