## Supplementary material for "Translatome analysis reveals microglia and astrocytes to be distinct regulators of inflammation in the hyperacute and acute phases after stroke": Table 1

| Dataset name | Reference | Filtering criteria used |
| --- | --- | --- |
| Disease-associated microglia cluster | (Keren-Shaul et al., 2017),  Table S3, mmc3 | FC > 1.5; p < 0.05 |
| Aged microglia cluster | (Hammond et al., 2019), Table S1, Cluster OA2 | FC > 1.5; FDR < 5E-88 |
| Demyelination-injury-responsive microglia cluster | (Hammond et al., 2019), Table S1, Cluster IR2 | FC > 1.5; FDR < 4E-106 |
| LPS-responsive vs. adult microglia | (Bennett et al., 2016), Dataset S01, Adult to LPS | FC > 1.5; FDR < 0.05 |
| 2 days post-TBI vs. sham microglia | (Izzy et al., 2019), Table S1 | FC > 1.5; q < 0.05 |
| 14 days post-TBI vs. sham microglia | (Izzy et al., 2019), Table S2 | FC > 1.5; q < 0.05 |
| 60 days post-TBI vs. sham microglia | (Izzy et al., 2019), Table S3 | FC > 1.5; q < 0.05 |
| End-stage vs. aged control microglia | (Chiu et al., 2013), Table S3 | FC > 1.5; FDR < 0.05 |
| LPS vs. PBS microglia | (Sousa et al., 2017), Data EV1 | FC > 1.5; FDR < 0.05 |
| 3 hour LPS-specific vs saline astrocytes | (Hasel et al., 2021), Table S2 | FC > 2; FDR < 0.05 |
| 24 hour LPS-specific  vs saline astrocytes | (Hasel et al., 2021), Table S2 | FC > 2; FDR < 0.05 |
| 72 hour LPS-specific vs saline astrocytes | (Hasel et al., 2021), Table S2 | FC > 2; FDR < 0.05 |
| 2 year old vs 10 week cortical astrocytes | (Clarke  et al., 2018), Dataset S3 | FC > 2; FDR < 0.05; FPKM ≥ 5 in 2-y-old samples for upregulated genes |
| EAE vs unimmunized astrocytes | (Borggrewe  et al., 2018), Table S5 | FC > 2; FDR < 0.05 |
| Alzheimer’s disease vs wild type astrocytes | (Orre et al., 2014), Table 1 | Top 50 upregulated genes, FDR < 0.05 |
| 3 day tMCAO vs sham astrocytes | (Rakers et al., 2018), Table S1 | FC > 1.5; FDR < 0.05 |
| 24 hour tMCAO vs sham astrocytes | (Zamanian et al., 2012), Table 1 | Top 50 upregulated genes, FDR < 0.05 |
| SOD1 vs wild type | (Sun et al., 2015), SI Table 4 | FC > 0.5; FDR < 0.05 |
| 2 year vs 4 month old visual and motor cortex astrocytes | (Boisvert et al., 2018), Table S3 | FC > 1.5; FDR < 0.05 |

**Table 1. Sources and criteria for microglia and astrocyte gene sets for Gene Set Enrichment Analysis (GSEA).**
